# GENIE: a multi-agent system for closed-loop autonomous microscopy

**DOI:** 10.64898/2026.10.12.758370

**Authors:** Wilson Nieves Vasquez, Jun Lin, Joshua Azukas, Evan Yang, Chi-Kuo Hu, Tengfei Ma, Lina Carlini

**Author notes:** Equal contribution (Jun Lin, Joshua Azukas).

## Abstract

Microscopy is foundational to biology, enabling direct observation of otherwise inaccessible cellular structures and processes. Artificial intelligence has advanced microscopy through acquisition control, image analysis, and natural language instrument operation. No system has unified these steps into a complete experimental loop. We introduce an autonomous multi-agent system that turns plain language goals into measurements and statistically tested answers, gating and executing every action in deterministic code. This sets the groundwork for extending microscopy from biological observation to autonomous investigation.

## Introduction

Spatial organization is central to cell biology and light microscopy makes this organization measurable. Extracting quantitative, biological information from raw image data requires a series of interconnected decisions, where acquisition and analysis inform one another. Observations at the microscope guide what to acquire next, while analysis determines whether the evidence is sufficient and how subsequent acquisitions should be adapted. Artificial intelligence has advanced individual steps in this process^1–4^. For instance, smart microscopy uses observations during acquisition to adapt subsequent imaging^5–9^, and language agents now assemble imaging pipelines on request^10^. Language models operate instruments with planning and hardware abstraction and return measurements for interpretation^10–17^. Agents write and run analysis workflows^18–20^. These systems automate individual stages, but the experimental loop, from a stated goal through acquisition to a statistically tested answer, has been closed in several fields^21–23^, but not in light microscopy^24,25^.

We introduce GENIE (Grounded Empirical Navigator of Instruments and Experiments), an autonomous multi-agent system that takes one plain language goal, acquires data and decides what the data supports. It closes the experimental loop by turning a stated goal into a statistically tested answer. Here, GENIE measured mitochondrial membrane potential in live cells and found that it is spatially coherent: the potential of a region predicts that of neighboring mitochondria over a coherence length of 20 to 30 µm. These findings reveal spatial organization of a fundamental bioenergetic property.

## Results

### Design and Implementation

GENIE is software that converts a question posed in natural language into a statistically defended answer. It carries an experiment autonomously from a goal, through acquisition, to quantitative analysis. The user specifies the scientific question, can provide experimental and analysis constraints, sets the operating permissions, and may intervene at any point. GENIE develops and executes the workflow, makes methodological and data-dependent decisions, and adapts its actions according to acquired data. It writes and revises the data-processing code and selects the statistical tests and uncertainty measures used to evaluate the results.

GENIE uses a large language model (LLM) for reasoning, vision models to interpret acquired images, and knowledge modules that combine stored facts with Bayesian estimates. Its controller cycle carries a user-provided goal through the following five stages.

1) Plan: The stated goal is first broken into an ordered set of steps, turning each step into concrete instructions and their parameters. 2) Gate: This stage screens each instruction against the permissions and physical limits that apply to it. 3) Execute: GENIE runs the instrument commands and analysis code that passed the gate. 4) Sense: It examines what came back, such as an acquired image, and decides what it represents, such as a subcellular structure of interest. 5) Summarize: This stage reduces those results to what the next cycle needs and stores what has been learned. The five-stage cycle is repeated until that step converges to the intended target, ultimately fulfilling the user-provided goal (Fig. 1a, b; Supplementary Notes 1 to 3). The deterministic gate keeps actuation authority outside the model. GENIE writes its own analysis during the run, with the instrument still under its control; a critic checks that code until it converges, screens it for scientific validity and audits it against the data it came from. We describe each stage, the design and the architecture in the supplement (Fig. S1; Supplementary Notes 1 to 3). We transferred GENIE from the microscope on which it was developed to an independent system, where it measured transfection efficiency from a user-stated goal, demonstrating its modularity (Fig. S2).

**Figure 1.**
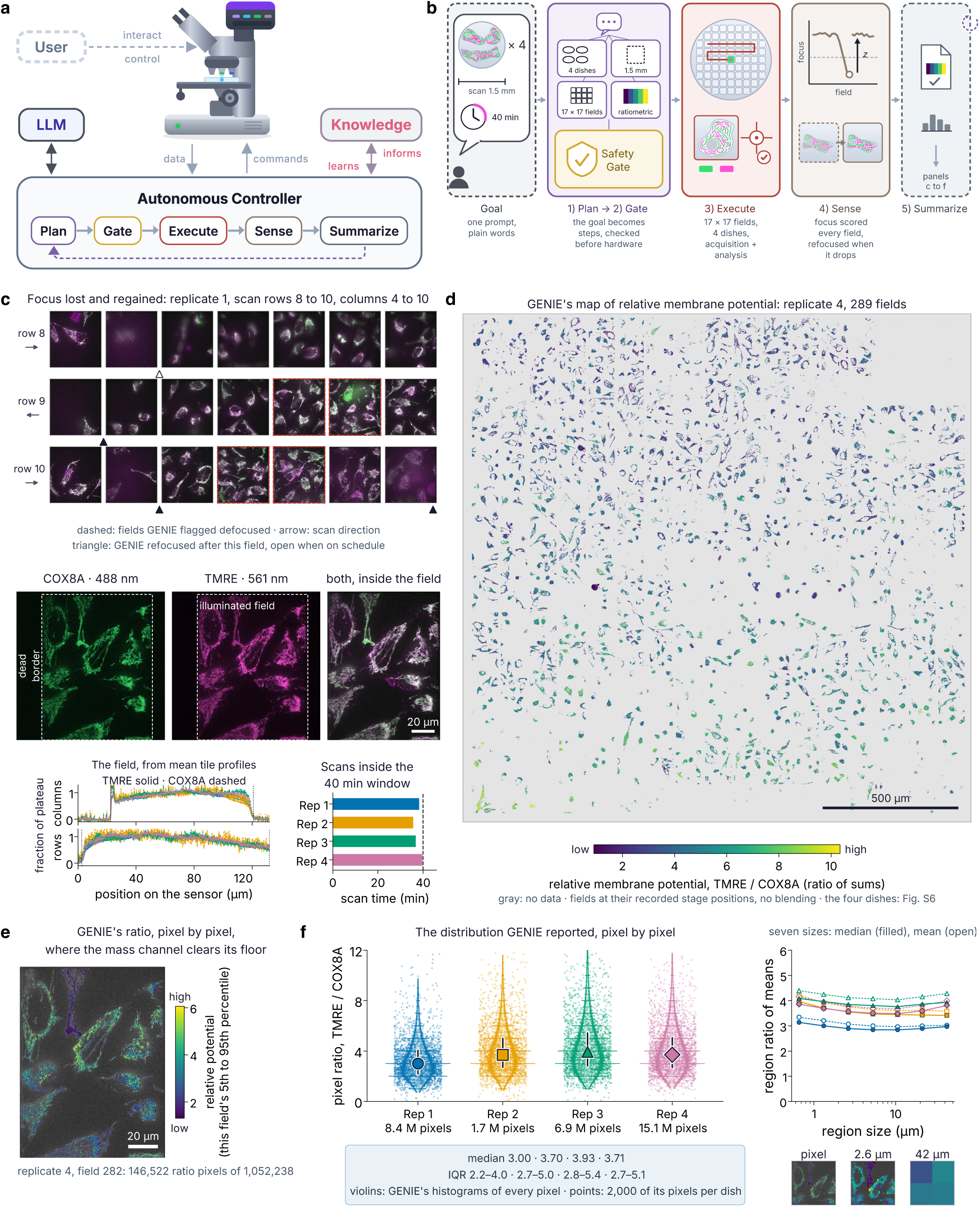
GENIE turns one plain language goal into a measured quantity. (a) A language model advises a controller of fixed code that runs plan (purple), gate (gold), execute (brick red), sense (brown) and summarize (slate blue), then returns to plan. The user and the language model are in dark gray. A hardware layer owns the devices, and a knowledge store (mauve) holds priors and findings (Fig. S1). (b) The run on that cycle, in the colors of a: the goal (the four dishes, the 1.5 mm scan and the 40-minute window); the plan and the safety gate it passes (four dishes, 1.5 mm, 17 by 17 fields, the ratiometric analysis); the scans, 17 by 17 fields per dish with the stage read back, TMRE then COX8A at every field; the focus score that triggers a refocus, sharpening the cells; and the summary that closes the loop and feeds c to f. Panels a and b are schematics drawn with AI assistance. (c) The events in the order they happened. At the top are three scan rows of replicate 1, with the fields GENIE flagged defocused (dashed) and its refocuses after a field (triangles; open, scheduled). In the middle is one field of replicate 4, with the dead border, the illuminated field GENIE set from the tiles (dashed), and both labels inside it. At the bottom are the mean tile profiles with the edges found (TMRE solid, COX8A dashed), and the four scans against the 40-minute window. (d) GENIE’s map of relative mitochondrial membrane potential for replicate 4, with fields at their recorded stage positions and gray where there is no ratio (all dishes, Fig. S6). The color runs from low (purple, 0.91) to high (yellow, 10.3), GENIE’s pooled 1st to 99th percentile over the four dishes, and the first scan row is at the bottom. (e) The ratio pixel by pixel over the COX8A image of the field in c, colored from low (1.3) to high (6.1), this field’s 5th to 95th percentile. (f) The distribution of each dish as violins from GENIE’s histograms of every pixel, with 2,000 of its pixels per dish as points and medians with the interquartile range. On the right are the median (filled) and mean (open) of the region ratio at the seven sizes, and under them an 84.5 µm square of the field of e (replicate 4, field 282) at three of those scales: the pixel, GENIE’s 2.6 µm regions (the smallest size of its whole-dish test) and its 42 µm regions (the largest), on the color range of e. Scale bars, 20 µm (c, e) and 500 µm (d).

**Figure 2.**
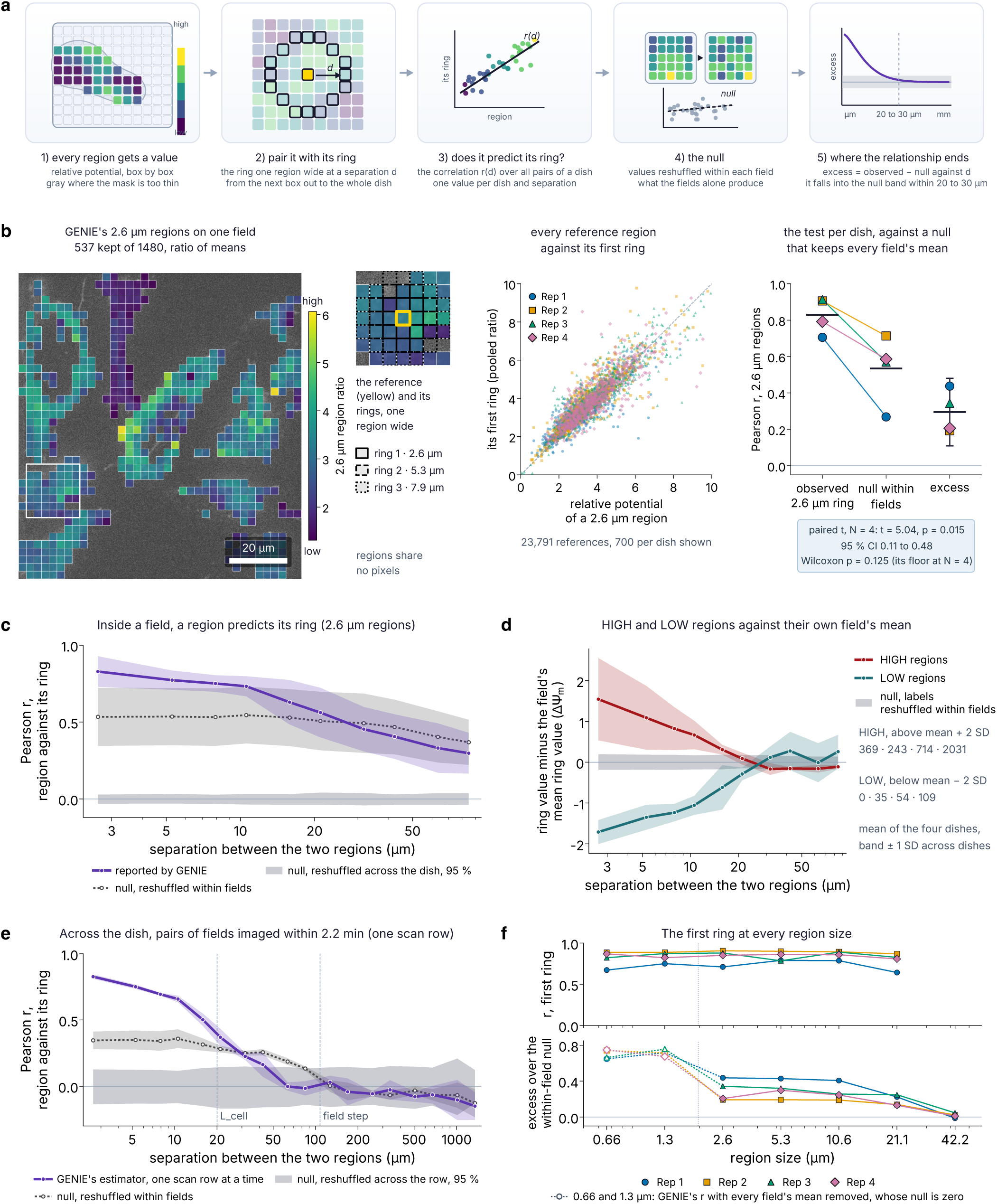
A region predicts its neighborhood, and the relationship ends within 20 to 30 µm. (a) The test runs from a region’s value to the excess of its correlation with its ring over the within-field null, followed with separation d. This schematic was drawn with AI assistance. Panels b to e use 2.6 µm regions; the other sizes are in Fig. S8. (b) Left, GENIE’s regions on one field of replicate 4, colored from low (1.3) to high (6.1), with the reference (yellow) and its first three rings. Middle, every reference region against the pooled ratio of its first ring (700 per dish shown). Right, the first ring per dish as observed correlation, within-field null and excess; the paired t on the excess gives N = 4, t = 5.04, p = 0.015. (c) The correlation against separation inside a field as reported by GENIE (purple) and its within-field null (gray dotted), over the 95 % band of the across dish null (gray). Both regions of a pair lie in the same field, so c uses every retained field of the whole scan and acquisition time cannot enter it; e pairs fields across the dish and is therefore restricted to one scan row. In c to e, lines are the mean of the four dishes and bands one SD across dishes. (d) The ring partners of HIGH regions (above the dish mean + 2 SD; red) and LOW regions (below the mean − 2 SD; teal) minus the mean ring value of the references in their own field, a difference on the scale of ΔΨm (a ratio with no unit), so that a field whose ΔΨm is high throughout does not inflate the offset; the band is the null with the class labels reshuffled within each field (gray, 200 reshuffles). HIGH regions sit above their field and LOW regions below it at the first rings, and both reach their field’s mean by about 30 µm. This baseline was computed by us from GENIE’s regions with GENIE’s rules; GENIE’s own curve against the dish mean is in Fig. S9b. (e) The sweep across the dish on pairs of fields imaged within about 2.2 min, one scan row at a time, with nothing corrected: GENIE’s estimator and its two nulls on every grid row of every dish (67 rows; 17, 16, 17 and 17), the median over rows per dish, then the mean of the four dishes (purple), its null reshuffled within fields (gray dotted) and its null reshuffled across the row (gray band, 95 %). A ring resting on fewer than 100 reference regions in a row is left out. The correlation is 0.83 at 2.6 µm against a null of 0.35, and the excess is gone by 32 µm. The guides mark L_cell, GENIE’s median area-equivalent cell diameter (19.8, 20.0, 20.0 and 21.2 µm in the four dishes), and one field step (107.6 µm). Within one row the time trend of the signal is removed by construction and nothing is corrected; over the whole scan the correlation stays above the null to about a millimeter, which is the decline of the TMRE signal during the 38 minutes of imaging and not the dish (Fig. S12). GENIE reported that long range correlation as the fields’ component and drew no conclusion from it. (f) The first ring at every region size from 0.66 to 42.2 µm, per dish: top, GENIE’s correlation between a region and its first ring; bottom, its excess over the within-field null, GENIE’s whole-dish record from 2.6 µm on. At 0.66 and 1.3 µm GENIE ran only its within-field sweep, whose correlation removes every field’s mean first and whose null is therefore zero; those two sizes are drawn dotted with open markers (0.65, 0.74, 0.65 and 0.75 at 0.66 µm; 0.72, 0.71, 0.74 and 0.67 at 1.3 µm).

### GENIE designs measurements and statistical analyses to investigate mitochondrial membrane potential

Mitochondrial membrane potential, a widely used indicator of mitochondrial function^26^, can vary several fold among mitochondria^27,28^. While spatial differences in mitochondrial membrane potential have been reported^27^, the organization of this heterogeneity across scales and the extent of local similarity remains unknown. To address this, we used GENIE to control a custom spinning disk confocal microscope. We asked it to map relative mitochondrial membrane potential in live cells expressing mStayGold-COX8A (hereafter COX8A), a reference signal for mitochondrial content, and stained with the potential-dependent dye TMRE (tetramethylrhodamine ethyl ester)^29–31^. Ratiometric analysis of these two signals normalizes TMRE fluorescence to the reference, a proxy for mitochondrial mass, and provides a relative readout of membrane potential. Hereafter, ΔΨ_m_ denotes this relative fluorescence readout. In the same prompt, we asked whether a region’s ΔΨ_m_ predicts that of other regions across spatial scales. We gave GENIE acquisition and analysis specifications, including the dye’s time window and the need for multiple biological samples. GENIE made the methodological choices needed to carry out these requirements (Fig. S3; the full prompt is in Supplementary Note 7).

Given the acquisition constraints, GENIE defined and executed the following imaging workflow. For each dish, GENIE selected a 17 × 17 grid (289 fields) with a recorded step size of 89.76 µm. As specified, it acquired the TMRE channel before the COX8A channel at each position and used brightfield to autofocus; in addition to the scheduled refocus every eight fields, it refocused whenever its self-defined focus metric dropped. This occurred 38, 20, 39 and 39 times in the four dishes (Fig. S4a, b; Methods, “GENIE’s control of acquisition”). The stage position was read back at every field, preserving spatial traceability for subsequent user inspection. This workflow was carried across all four dishes, pausing for user-mediated sample exchange and resuming acquisition when each new dish was loaded. The four scans took 38.2, 35.6, 36.7, and 39.7 min, respectively, each within the user-specified 40-min acquisition window.

Next, GENIE coded the steps needed to convert the raw fluorescence data into a ΔΨ_m_ measurement. It determined the usable illuminated region from the mean intensity profiles of 2,296 acquired tiles and fixed it as a single crop region for analysis (Fig. S5a). For each image, it subtracted background and generated a mitochondrial mask by thresholding the COX8A (mass proxy) channel. Within these masks, GENIE computed ΔΨ_m_ per pixel as the ratio of the TMRE intensity to the COX8A intensity and assembled the fields into one map per dish at their recorded stage positions (Fig. 1d, e; Fig. S6). In every dish, the distribution was unimodal and right skewed (Fig. 1f; Fig. S7). GENIE also performed a region-based version of this analysis, dividing each image into a grid of square boxes from 0.66 to 42.2 µm on a side; a box with at least 25 masked pixels is a region (Fig. 1f; Methods, “Regions”). Within each region, GENIE estimated ΔΨ_m_ as the ratio of the mean masked-pixel intensities in the TMRE and COX8A channels; the median across the regions pooled from all images varied by ∼10 % across region sizes (Fig. 1f). GENIE further defined a rule for eroding the mask at its boundary, but one of the independent verifications found an inconsistency in how its erosion step applied that rule, one that does not affect the downstream results (Fig. S5b, c; Methods; Supplementary Note 5). Together, these data suggest that ΔΨ_m_ is a characteristic property of these cells, consistent from the pixel scale to regions 42 µm across.

Using the same multiscale regions, GENIE measured the distance over which a region’s ΔΨ_m_ remained detectably correlated with that of its surroundings, which we call the spatial coherence length (Methods, “Spatial correlation and its nulls”). It calculated the Pearson correlation between each region and successive surrounding rings. To determine whether the measured correlation exceeds that expected by chance, GENIE shuffled region values 200 times within single fields of view (hereafter, within-field null) and across the reconstructed dish. It also chose the statistical tests and uncertainty measures for the four dishes (Methods, “Spatial correlation and its nulls”). We selected 2.6 µm regions (24 pixels on a side) for reporting because this was the smallest size for which GENIE completed both the correlation analysis across the reconstructed dish and its within-field null computation. This size allows local sampling of mitochondrial signal while limiting spatial averaging across neighboring cells. We assessed the robustness of the spatial findings using GENIE’s analyses at other region sizes: regardless of the size, regions were strongly correlated beyond the within-field null and the correlation declined with distance, falling significantly by ∼20 µm (Fig. 2b, c, f; Fig. S8b, c). The excess correlation over the null ended by 30 to 40 µm at every size, a coherence length on the order of a cell diameter (Fig. S8c). High- and low-potential regions, classified by their deviation from the mean ΔΨ_m_, had neighbors with correspondingly high and low ΔΨ_m_ relative to their in-field mean ring value. Both deviations diminished with distance, suggesting local physical clustering of mitochondria with similarly elevated or reduced mitochondrial potential (Fig. 2d; Fig. S9b).

Extending the analysis across the dish, GENIE’s whole-dish correlations remained above the within-field null over hundreds of micrometers. Due to the serpentine stage-scan pattern, a field’s position across the reconstructed dish covaried with its acquisition time (Fig. 1d; Fig. S6). GENIE also reported a decline in ΔΨ_m_ over acquisition time, creating a possible confound between temporal and spatial variation. GENIE considered the within-field null sufficient to account for this temporal effect and concluded that correlations remained distinguishable from the null at separations up to ∼30 µm (Fig. S11f). Restricting the same sweep to pairs of fields imaged within about 2.2 min, one scan row at a time, removes the temporal effect by construction: the excess over the within-field null was 0.48 at 2.6 µm and ended by 32 µm (Fig. 2e; Fig. S12a, b). The GENIE-independent verifications (Supplementary Note 5) confirmed the local correlation within 20 to 30 µm (Fig. S10a, g; Fig. S11d–g) and, in agreement with GENIE, showed that the apparent long-range correlation was indistinguishable from the temporal effect introduced by the scan (Fig. S12c, d). The within-field sweep avoids this temporal effect because both regions in each pair lie in the same field. This analysis yields the same coherence length (Fig. 2c; Fig. S8c). Together, GENIE’s analyses indicate that ΔΨ_m_ is coherent over 20 to 30 µm.

## Discussion

GENIE is a multi-agent system that takes a plain-language goal, acquires data, decides what the data supports and returns a statistically supported answer. Here it closed the loop: GENIE planned and acquired the scans, then wrote, checked and ran its analysis on four samples. From a user-posed question, GENIE found that ΔΨ_m_ is coherent over a length of 20 to 30 µm. This spatial organization is consistent with prior observations of coordinated mitochondrial behavior^32–35^ and supports a model wherein mitochondrial states are coupled within and, possibly, across cells. Future work will be required to determine the mechanisms underlying this coherence.

This run also illustrates the importance of critical checking and human oversight. GENIE measured a temporal dependence in ΔΨ_m_ and used a within-field null that was sufficient to establish the local correlation. We verified this with independent analyses and subsequently showed that the apparent long-range correlation was indistinguishable from that arising from the temporal dependence (Fig. S12c, d). The local correlation within 20 to 30 µm, however, persisted after the measured temporal trend was accounted for. Independent verification also identified an inconsistency in GENIE’s application of its self-defined mask-erosion rule (Fig. S5b, c). Neither issue changed the biological conclusion, but both illustrate the importance of provenance and independent verification in evaluating autonomous analyses.

Future work will extend GENIE from answering predefined questions to choosing measurements that discriminate among hypotheses. For quantitative microscopy, this will shift researcher effort from implementing acquisitions and analyses toward formulating scientific questions and developing new ways to answer them.

## Methods

### Implementation of GENIE

GENIE is a Python application built around a controller written as fixed code and advised by a language model; its architecture, the roles of the language model and its knowledge stores are described in Supplementary Notes 1 to 3 and Fig. S1. All experiments used GENIE, running under Python. The live-cell experiment (Figs. 1 and 2) used Anthropic’s Claude Opus 4.6 for the acquisition and Claude Fable 5.1 for the analysis. The experiment on the second microscope (Fig. S2) used Claude Fable 5.1 only. Only text reached the models: the goal, the state of the experiment, the generated code and its output. Raw image data was processed locally and not transmitted to the language models.

For the experiments described here, the controller cycle (plan, gate, execute, sense, summarize) ran once per subgoal at four of GENIE’s seven autonomy levels. Every proposed hardware action was screened by the deterministic gate, which enforced the illumination powers and exposures fixed by the prompt, the stage motion limits and the identification of every device before it was written to. The gate made no language model call. GENIE drove the microscope through its hardware layer, which identified the connected devices at start up and, after each command, read the device back and recorded both the requested and the applied state with the frame. For the live-cell experiment (Figs. 1 and 2), acquisition of all four dishes was completed before analysis. The complete mission directory was then copied to a separate computer without an attached microscope, where analysis continued under the original goal from the saved records.

GENIE screened the code it generated for unsafe operations, used its critic to check scientific validity and audited the code against its execution output (Supplementary Note 2). The critic’s statistical rubric (Supplementary Note 2) required statistical analyses that treated independent experimental replicates as the units of comparison, effect sizes with confidence intervals and stated assumptions. During the live-cell analysis, GENIE’s watcher agent, a language model agent that follows the run, supplied guidance to the planner and assisted with script revision. Its messages were logged as standing constraints that guided subsequent planning and analysis (Supplementary Note 2).

The run was recorded in an event log, a per-decision audit trail and a provenance record that labeled each raw file by sample, replicate and channel. Every numerical constant had to come from the user, the hardware, file metadata or a declared prior, or be reported as unknown; we refer to this rule as constant sourcing. GENIE read the pixel size, 0.11 µm per pixel, from the tile metadata, a configured value that the operator confirmed. The deliberation engine required the corresponding deliverables to exist before declaring a phase complete, and rejected outputs were quarantined rather than deleted.

Key scientific dependencies were numpy, scipy, scikit-image, scikit-learn, pandas, OpenCV, tifffile, matplotlib, PyTorch, torchvision, Cellpose^1^, and Segment Anything^36^. GENIE can also operate a microscope through Micro-Manager^37^: its hardware layer has an optional back end that works through Pycro-Manager^38^. Micro-Manager was not used in this work; GENIE drove both microscopes directly through its own hardware layer. GENIE’s architecture, the roles of the language model and its knowledge stores are described in Supplementary Notes 1 to 3 and Fig. S1.

### Microscopes and imaging hardware

Live cell imaging (Figs. 1 and 2) was performed on a Nikon Ti inverted microscope with Perfect Focus, a Prior motorized XY stage and a Yokogawa CSU-X1 spinning disk confocal unit. Samples were imaged through a Nikon 100x oil immersion objective (NA 1.4) onto a Teledyne Photometrics Prime 95B sCMOS camera controlled through PVCAM (1,200 by 1,200 pixel frames, 0.11 µm per pixel). Excitation came from a Vortran Laser Technology laser launch with 405, 488, 561 and 640 nm lines, each switched by TTL and set in power by an analog voltage from a TriggerScope controller (Advanced Research Consulting). TMRE was excited at 561 nm (9 ± 2 mW, measured at the output of the delivery fiber) and imaged through a 629/62 emission filter (the TMRE channel), and mStayGold-COX8A was excited at 488 nm (14 ± 2 mW at the fiber output) and imaged through a 520/35 emission filter (the COX8A channel). All laser powers in this paper were measured at the output of the fiber that delivers the laser light to the microscope, at the power setting used for imaging, not at the sample. Both filters sat in the CSU-X1 filter wheel, and the Ti filter turret was at its empty position. Exposure was 50 ms per channel. The two channels were acquired in sequence at each field (TMRE first) without moving the stage, and autofocus ran on brightfield images every eight fields. GENIE drove every device directly through its hardware layer, without Micro-Manager, and each tile’s header records the settings sent to each device.

The fixed cell experiment (Fig. S2) was performed on a second microscope, a Nikon Ti inverted microscope with a Yokogawa CSU-X1 spinning disk confocal unit, through a Nikon Plan Apo VC 100x oil immersion objective (NA 1.40) onto a Hamamatsu ORCA-Flash4.0 V2 sCMOS camera (C11440-22CU; 2,048 by 2,048 pixels, 6.5 µm pixels, 0.065 µm per pixel). The XY stage and the light path were driven through the Ti. Excitation came from a six-line Vortran Laser Technology laser launch (405, 445, 488, 505, 561 and 640 nm) controlled over a serial interface compatible with the Lumencor Spectra. DAPI was excited at 405 nm (60 mW at the fiber output) and imaged through a 460/50 emission filter, and H2B-mRuby was excited at 561 nm (30 mW at the fiber output) and imaged through a 488/561 dual band emission filter, both in an ASI FW-1000 filter wheel. Exposure was 400 ms per channel. GENIE ran this microscope without Micro-Manager or the microscope’s own acquisition software, using the same hardware layer and a configuration built from the microscope’s own exported settings.

### Cell culture and live-cell labeling

HeLa TRex cells were cultured in DMEM (Cat 11995073; Gibco) supplemented with 10 % FBS (Cat FB73; Alkali Scientific) and incubated in 5 % CO_2_ at 37 °C. To generate a stable HeLa TRex cell line expressing mStayGold-COX8A, cells were co-transfected with 2 µg of qTAG-C-Solo-mStayGold-COX8A (Addgene plasmid #227309) and 2 µg of pX330-PITCh-COX8A (Addgene plasmid #227308) using Lipofectamine 3000 and P3000 reagents (Cat L3000008; Thermo Fisher Scientific), following the procedure described by Philip et al.^39^. After 24 h post-transfection, the culture medium was replaced with fresh medium, and cells were incubated at 37 °C with 5 % CO_2_ until 72 h post-transfection. Cells were subsequently isolated by fluorescence-activated cell sorting using a BD FACSAria IIIu Cell Sorter (BD Biosciences) equipped with a 488 nm excitation line and a GFP emission filter. For live-cell imaging, mStayGold-COX8A cells were loaded with 25 nM TMRE, the potential-dependent dye imaged at 561 nm, and imaged live at 37 °C in L-15 medium; TMRE was imaged within 40 minutes of the first exposure, the imaging window we use for the dye based on reported protocols^30,31^.

### Transient transfection and fixation

For the fixed-cell experiment, HeLa cells were plated onto 22 mm #1.5 coverslips (Cat 50-233-5828; Fisherbrand) and transfected with 500 ng of pKanCMV-mRuby3-10aa-H2B (Addgene plasmid #74258) with Lipofectamine 3000 according to the manufacturer’s protocol. At 24 h post-transfection, cells were fixed with 4 % paraformaldehyde (Cat 15710; Electron Microscopy Sciences) at 37 °C with 5 % CO_2_ for 10 minutes, rinsed three times with PBS, mounted with VECTASHIELD mounting medium with DAPI (H-1200-10; Vector Laboratories) and sealed with nail polish.

### GENIE’s control of acquisition

The four biological replicates were separate dishes from independent cultures, loaded one at a time, with GENIE pausing after each to request the next. Since the sample was live and TMRE is imaged within a limited window after loading, the prompt fixed the illumination at 10 % and 15 % of the laser output for 561 nm (TMRE) and 488 nm (COX8A) (9 mW and 14 mW at the fiber output; 50 ms exposure each), told GENIE that the dye was good for 40 minutes from the first exposure, and required acquisition to start at once, before any pilot image or analysis. The prompt named the two channels and their roles, the 1.5 mm by 1.5 mm region and the replicate count, described what the analysis had to respect, and left every other value to GENIE; it did not name the fluorophores. GENIE’s grid scan module tiled the region into a 17 by 17 raster of 289 fields per dish at a recorded step of 89.76 µm, acquiring the TMRE channel then the COX8A channel at each position before moving and revisiting no field, with autofocus on brightfield every eight fields and whenever its focus metric fell below threshold, and the stage position read back at every field, within 0.9 µm of the commanded position. The self-defined focus metric is the mean-normalized variance of the Sobel gradient magnitude over the central half of each acquired TMRE tile, compared with the median of the tiles acquired since the last refocus; a refocus was triggered when it fell below 0.55 of that median. The refocus itself searched z on brightfield frames, scoring each by the same gradient metric within the cells that a segmentation model (Cellpose) found in it. The four scans took 38.2, 35.6, 36.7 and 39.7 minutes from the first to the last exposure, no pixel saturated in either channel, and the focal plane moved by 24, 57, 14 and 20 µm over a scan and was recovered at each autofocus event (Fig. S4b). The last eight fields of dish 2 were not acquired, and the scan record was rebuilt from the tile headers; 1,148 fields, 2,296 tiles, entered the analysis. Both channels of a field share one stage position, so no registration between them was needed.

### GENIE-executed analysis

#### Relative membrane potential (Fig. 1)

##### Illuminated field

The camera sensor is larger than the area the lasers illuminate, so every frame has a dark margin in which the sensor records only its own noise; we call this the dead border. GENIE measured the illuminated area from the data. For each channel and each dish, 8 groups in total, GENIE accumulated the mean row and column profiles over all 2,296 tiles. Along each profile it set an edge where the profile left the dead level by five standard deviations of the dead border. The TMRE channel gave four estimates of the illuminated rectangle, one per dish, and GENIE took their innermost envelope. It then checked the four COX8A estimates against that rectangle, with a tolerance of 15 pixels. The rectangle is rows 23 to 1199 and columns 205 to 1098 of the sensor (894 by 1177 pixels, 98.3 by 129.5 µm). GENIE saved it as an instrument constant. Every later step reads this constant and crops each tile to it (Fig. 1c; Fig. S5a).

##### Placement

Along the stage’s Y axis, the axis along which the stage stepped between scan rows, the illuminated field is 1177 pixels long but the step was only 978 pixels, so each field overlaps the next one in the scan by 199 pixels and the two share that strip of content. GENIE measured the shift of that shared content by cross-correlation on 15 pairs of fields whose estimates agreed: one recorded stage step of 89.76 µm corresponds to 978 pixels of image. The recorded stage positions were therefore short by a factor of 1.1985. GENIE applied that factor when it placed every field of a dish in one common frame. The same correlation showed that image rows run along the Y axis of the stage. Along X, fields in neighboring columns share no content, so there was nothing to test, and GENIE took the stage’s own sign for that axis. To count nothing twice, each field was given a window of 978 by 894 pixels. A region (Fig. 2a) belongs to the field whose window holds its center, and to that field only. The illuminated area outside the window is still used: it supplies the surroundings of regions near the edge of the window. For a reference region and its ring (defined below under “Spatial correlation and its nulls”), distances inside one field are exact. Distances between fields carry the placement error, 0.97 µm: the scatter of the measured shift between neighboring fields about the corrected positions, which remains after the correction above. GENIE reported this error with its placement records; being a random residual, it was not corrected further. It cannot enter the within-field sweep (defined below), which uses only distances inside one field. In the whole-dish sweep it affects only pairs whose members lie in different fields, and it is smaller than one ring width at the sizes reported (2.6 µm), so it can move such a pair into a neighboring ring but cannot create a correlation. At the first ring the two sweeps agree within 0.06.

##### Background, mask and erosion

GENIE estimated the background of each tile as the mode of its positive counts. It estimated the noise of each tile as the larger of two values: the standard deviation of the right flank of the count histogram, and the read noise it had measured from the dark frames, 7.2 ADU. It set a floor for each tile at the background plus three noise standard deviations. The median floors were 24.6 to 25.9 ADU for COX8A and 35.0 to 49.2 ADU for TMRE. The mask is the set of pixels of the COX8A channel above its floor, kept in connected components of at least 5 pixels. The same mask was applied to both channels, and the two channels were divided by one another to yield per-pixel ratiometric images of TMRE:mStayGold-COX8A (Fig. 1e). Next, GENIE swept an erosion of the mask from 0 to 10 pixels (0 to 1.1 µm) and chose the depth by a rule with two conditions, both to hold in every dish: 1) at least half of the mask pixels kept, and 2) a change of less than 2 % in the dish’s median field ratio. The ratio of a field is the mean TMRE count over its masked pixels divided by the mean COX8A count over the same pixels; the median is taken over the fields of the dish. One erosion iteration kept 50, 45, 51 and 53 % of the mask pixels in the four dishes, so no depth from 0.11 µm (one pixel) on met the first condition, and GENIE operated at a depth of 0. The recomputation (Supplementary Note 5) found why one iteration removed so much: GENIE’s erosion used a 3 by 3 square, which removes diagonal boundary pixels as well, so one iteration removes more than one pixel of depth. Eroded by Euclidean distance instead, a depth of 0.11 µm (one pixel) keeps 55 to 62 % of the mask and would have met the rule (Fig. S5b, c). Fields were retained for subsequent analysis by a focus rule on the COX8A image: its normalized Laplacian variance had to lie within three robust standard deviations of the dish median. This retained 283, 271, 289 and 289 fields, of which 277, 151, 267 and 288 carry a mask; dish 2 is the dim one, which is why only 151 of its 271 retained fields carry a mask.

##### Regions

A region is a square of 0.66, 1.3, 2.6, 5.3, 10.6, 21.1 or 42.2 µm on a side (6, 12, 24, 48, 96, 192 or 384 pixels; Figs. 1f and 2a), on a grid anchored at the origin of the window. Regions of one size share no pixels. The value of a region is its ratio of means: the mean background-removed TMRE count over its masked pixels divided by the mean COX8A count over the same pixels,

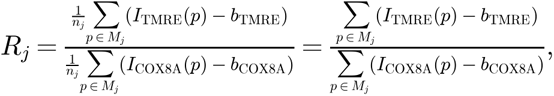

where *I*_TMRE_(*p*) and *I*_COX8A_(*p*) are the counts of pixel *p* in the TMRE and COX8A channels, *b*_TMRE_ and *b*_COX8A_ the background of the tile in each channel, *M*_j_ the masked pixels of region *j* (the mask within its box) and *n*_j_ their number; the 1/*n*_j_ cancels, so the ratio of means equals the ratio of the sums. Dividing pixel by pixel first and then taking the mean or the median of the pixel ratios,

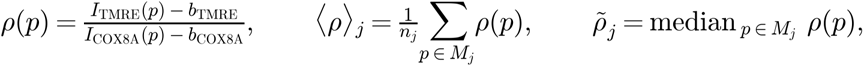

gives the same results: with GENIE’s own sweeps and nulls rerun with only the region value changed, the first-ring excess is 0.29 under all three estimators (p = 0.015, 0.015 and 0.022) and the mean curve meets the within-field null at 32 µm under all three (Supplementary Note 5; Fig. S13).

A region is kept when it holds at least 25 masked pixels. GENIE derived this floor from the noise bound of the denominator: every mask pixel’s background-removed COX8A count exceeds three noise standard deviations σ, so the sum over the *n* masked pixels exceeds 3*n*σ while its noise is about √*n* σ, and the relative noise of the denominator is therefore below

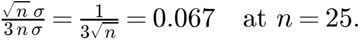

Regions below the floor stay in the tables, marked as excluded. GENIE also measured the resolution element of its mask as the median cross-sectional width of the mitochondria in the mask, their ridge width: 2 pixels, or 0.22 µm. It treated separations shorter than twice that width as a “blur” regime. Its smallest region, 0.66 µm, is three times that width. Every result was repeated across the seven sizes. The main text and the figures report the 2.6 µm size (24 pixels, 2.64 µm exactly). At that size the kept regions on the retained fields numbered 32,515, 6,470, 26,741 and 56,889.

##### Mitochondrial membrane potential distributions and classes

For each dish and each size, GENIE reported the histogram and the moments of the region ratio, together with a dip test of unimodality. The test found every dish unimodal at every size (p ≥ 0.92 at 2.6 µm). At 2.6 µm the means were 3.07, 3.75, 4.12 and 3.79, with standard deviations of 1.74, 1.54, 1.93 and 1.50, and the medians were 2.89, 3.55, 3.85 and 3.57. Every distribution was right skewed (Fig. S7). GENIE defined several classes: HIGH regions as more than two standard deviations above the mean and LOW regions as more than two below. At 2.6 µm this gave 369, 243, 714 and 2,031 HIGH regions (1.1, 3.8, 2.7 and 3.6 %) and 0, 35, 54 and 109 LOW regions. Figure 2d follows the surrounding rings of these classes (rings are defined under “Spatial correlation and its nulls”). The ring value is taken for each HIGH or LOW reference. The mean ring value of all references in its own field is subtracted from it, so that a field whose ΔΨ_m_ is high throughout does not inflate the offset. The HIGH curve plots the mean of these differences over the HIGH references. The LOW curve does the same over the LOW references. The value plotted is on the same scale as ΔΨ_m_, a ratio of counts with no unit. The curves are averaged over the four dishes. The null band is the same quantity after the class labels were reshuffled within each field (200 reshuffles). GENIE’s own version, against the dish mean, is Fig. S9b. GENIE also swept the multiplier from 1 to 3 standard deviations. HIGH regions fall from 7.2 to 14.0 % of all regions at one standard deviation to 0.3 to 1.5 % at three. LOW regions are present in every dish at one standard deviation (3.3 to 12.5 %) and in none at 2.5. Plots follow the SuperPlot convention^40^: per-region data are colored by dish, and per-dish summaries are overlaid. All inferential claims were made at the replicate level, as the prompt required: the biological replicate is the unit of inference, with N = 4.

#### Spatial correlation and its nulls (Fig. 2)

GENIE asked whether a region’s ΔΨ_m_ predicts that of its surroundings. Each region in turn served as the reference. GENIE grouped the other regions of the same size by their distance from the reference. The eight regions that touch the reference form ring 1. The regions about two region widths away form ring 2, and so on outward. Each ring is one region wide: the ring at separation m holds the regions whose centers lie between m – 0.5 and m + 0.5 region widths away (Fig. S13a). There are 16 separations, from one region width to 256 region widths (0.66 µm to 1.35 mm). The regions in a ring are its ring partners. The value of a ring is the ratio of means over all masked pixels of its partners, so a ring is measured the same way as a region. For each separation GENIE reported three things: the Pearson correlation between the reference values and the ring values, the slope of a linear fit of ring value on reference value, and the number of regions and masked pixels in each ring. At each separation, GENIE reported the mean and standard deviation across the four dishes for both the Pearson correlation coefficient and the fitted slope, with a 95 % confidence interval for each mean from the t distribution.

GENIE performed a spatial correlation analysis in two ways. First, it performed a within-field sweep, where each region was paired with the rings that fit inside its own field. Here, distances are exact, and GENIE pooled pairs of all retained fields within a dish. Second, a whole-dish sweep was performed. GENIE selected 6,000 reference regions for each dish and region size. These regions were paired with rings sampled from all fields in the dish. All fields were placed in a common coordinate frame, and distances between regions in different fields were calculated from the corrected stage positions.

Each sweep was compared with its corresponding null model. For the within-field sweep, values were randomly shuffled within each field. This disrupted spatial relationships within fields while preserving each field’s value distribution and mean. For the whole-dish sweep, values were shuffled across all fields in the dish, so differences between fields were no longer preserved. Each procedure was repeated 200 times, with the mask, region positions, and rings kept fixed. Excess correlation was defined as the observed correlation minus the within-field null correlation and represented spatial structure below the field scale. The field component was defined as the within-field null correlation minus the whole-dish null correlation and was not interpreted as a property of the sample. The coherence length of each dish, which GENIE’s records call the reach, is the distance beyond which the observed correlation remains within the 95 % band obtained from within-field shuffling.

To follow the whole range of the reference, GENIE split the references into fifths of the dish distribution. For each fifth it reported the mean value of the ring partners minus the mean over all references. It did the same for the HIGH and LOW classes, against a null that reassigns the reference labels while every reference keeps its ring; this null asks only whether the level of a reference carries information about its ring. GENIE also traced mitochondria, as connected components of the mask, and cells, as clusters of mitochondria around one nuclear void, and repeated the test with them as the unit; those results are not reported here.

The biological replicate was the unit of inference, with N = 4. GENIE ran a paired t test on the four per-dish excesses at the first ring to test whether the observed correlation differed from the within-field null. It reported the mean difference with its 95 % confidence interval, and alongside it the exact Wilcoxon signed-rank test and the sign test. For 2.6 µm regions the mean excess was positive and significant by the paired t test (Fig. 2b, right). The two rank tests gave p = 0.125. That is not a weaker verdict on the data: it is the smallest p value either test can give at N = 4, reached only when all four differences have the same sign, which they did. With four dishes there are only 16 ways the signs of the differences can fall, so a test that looks only at signs and ranks can never get below p = 0.125. The t test is not limited in this way because it also weighs how large the differences are. GENIE stated both limits itself: normality cannot be assessed at N = 4, and the rank tests cannot reach significance at this N. The claim therefore rests on the t test, with its confidence interval, and on the agreement in sign of all four dishes.

For 2.6 µm regions, the within-field correlation at the first ring was 0.71, 0.91, 0.88 and 0.85 in the four dishes, with slopes of 0.58, 0.86, 0.80 and 0.77. Over all pairs of a dish, the correlation at the first ring was 0.71, 0.91, 0.91 and 0.79. The within-field null there was 0.27, 0.71, 0.57 and 0.59. The excess was therefore 0.44, 0.19, 0.34 and 0.21: mean 0.29, 95 % CI 0.11 to 0.48, paired t = 5.04, p = 0.015, Wilcoxon p = 0.125 (Fig. 2b). The field component itself was 0.27, 0.71, 0.57 and 0.59, mean 0.53. The excess fell to 0.11, 0.03, 0.09 and 0.00 at 21.1 µm, and from 31.7 µm on it lay within 0.04 of zero or below it. The coherence length was 21.1, 31.7, 31.7 and 31.7 µm. GENIE ran its own replicate-level inference at 5.3 µm (48 pixels), the size it reported, where the excess was 0.43, 0.19, 0.32 and 0.30 (t = 6.41, p = 0.0077). The test at 2.6 µm applies the same rule to its 2.6 µm values. Software: numpy, scipy, scikit-image, pandas, tifffile, matplotlib, PyTorch and diptest.

#### Verification of GENIE’s results

All results were verified with GENIE-independent methods. The images were inspected for quality during and after each acquisition, and the quantitative checks were then performed by the authors without GENIE: an expert verification of the image analysis and a recomputation from the raw tiles, both described in Supplementary Note 5.

#### Use of Artificial Intelligence

GENIE used Anthropic’s Claude Opus 4.6 for acquisition and Claude Fable 5.1 for analysis; the second-microscope experiment used Claude Fable 5.1. The authors designed the experiments, wrote the independent verification analyses scripts with AI assistance, interpreted the results and wrote the manuscript.

Separately, the authors used Claude Fable 5.1 through Claude Code to assist with code development for inspection of mission logs, verification analyses and figure preparation, and with manuscript formatting, checks against the recorded outputs and preparation of submission files. The authors designed and specified these tasks and reviewed the resulting code and outputs. Schematics were created as code-generated vector drawings; no image-generating model was used. The authors take responsibility for the analyses, figures and manuscript.

## Data availability

The raw images with their stage positions, the numerical results of the analysis (per field, region and cell values, the separation sweeps with their nulls, and the source data of every figure) and the tables of the independent verifications are deposited in Zenodo (https://doi.org/10.5281/zenodo.23245979) and publicly available. The GENIE software is not included (see Code availability).

## Code availability

The GENIE code is not released with this preprint.

## Acknowledgments

This work was supported by the Laufer Center for Physical and Quantitative Biology and the AI3 Institute (award #106416 to L.C. and T.M.). W.N.V. and J.A. were supported by the National Institute of General Medical Sciences of the National Institutes of Health (award K12GM102778). C.-K.H. was supported by the National Institute on Aging of the National Institutes of Health (award DP2 AG077431). T.M. was supported by the National Science Foundation (grant 2514002) and the Patient-Centered Outcomes Research Institute (award 102727). The content is solely the responsibility of the authors and does not necessarily represent the official views of the National Institutes of Health. We thank Thomas Geer (Nobska) for microscope assistance, Benjamin Lin for providing the H2B-mRuby3 plasmid (https://www.addgene.org/74258/), Todd Rueb (Research Flow Cytometry Core Facility, Department of Pathology, Stony Brook Medicine) for technical support with the FACS ARIA, Alexandria Lewkowski for help with the schematic panels and pilot experiments, Nathaniel Sweet for help in getting started on the microscope in C.-K.H.’s laboratory, and Dora Mahecic for comments on the manuscript.

## Author contributions

W.N.V. and L.C. conceived GENIE. W.N.V. wrote the software. L.C. designed the experiment. W.N.V. performed experiments, prepared the figures and performed the recomputation of GENIE’s outputs from the raw tiles (Supplementary Note 5). J.L. prepared all samples, generated the cell line and performed experiments. J.A. performed the expert verification of GENIE’s analysis. T.M. and E.Y. performed an independent verification of GENIE’s architecture and code implementation. W.N.V., L.C., T.M. and E.Y. wrote the manuscript with input from all authors. W.N.V., J.A., T.M., C.-K.H. and L.C. supervised the work and acquired funding. All authors read and approved the final manuscript.

## Competing interests

W.N.V. and L.C. are inventors on a pending patent application covering GENIE. The GENIE software is copyright 2026 Research Foundation for SUNY and is not released with this manuscript. The authors declare no other competing interests.

## Supplementary Information

### Supplementary Note 1: On Architecture

#### Plan

An LLM decomposes the goal into an ordered set of phases. A router system then expands these into concrete actions and parameters. Once prompted, the planner proceeds on its own, and the user can interrupt it at any point. The planner holds no execution privileges to any device.

#### Gate

GENIE screens every instruction before it reaches an instrument. This stage is done entirely by fixed code, with no language model involved, so its checks can be enumerated. Which checks apply depends on what is being attempted, on what sample, and with which devices. Checks can include: an authorized permission tier set by the user, motion bounds, per capture illumination limits for that sample type, whether the target device has been identified and is still answering, the directories a generated script may touch, etc. GENIE may veto an instruction, or perform further screens. Acquisition constants come from the instrument itself, read back where available (camera readout time, sensor dimensions, bit depth and the like); users can also configure them.

#### Execute

Two paths exist under this stage. Hardware instructions go to GENIE’s hardware layer, a distinct part of the system that owns all contact with devices. At start up, GENIE enumerates what is connected and tests each capability a device claims, keeping only what is verified, then binds the verified devices to specific roles and speaks to each through its native interface. Every hardware instruction is confirmed by reading the instrument back, the stage position, the device states and the returned frame, before it is reported as done; no action is claimed as performed without that read back. The code GENIE writes for itself takes the other path and passes more screens. A language model writes any script, which needs to pass several checks for unsandboxed code and efficient computation patterns. A rejected script is regenerated with its violations attached and checked again, and the exchange is abandoned if it stops converging. That write, screen and regenerate exchange is a generator and critic loop. A watcher agent, a second language model instance that follows the run, supplies hints to the planner and helps a script pass the critic; its hints enter the mission as standing constraints and are logged like the user’s messages. Surviving scripts are reviewed for cost and scientific validity and audited against their own output for fabricated results (Supplementary Note 2). Screening is therefore a series of gates spread across the cycle, differing in what they examine, how harsh they are, and whether they can stop the run. All of it runs inside the cycle that drives the instrument.

#### Sense

This is where GENIE decides what it is looking at, and the stage into which external models plug. Cellpose, Segment Anything and task specific networks each register a description of what they do: task, imaging domain, suitable channels, GPU need, runtime. The vision model for a step is chosen against its description at run time, so any registered vision model can be used. Users can also add custom vision models here. Frames are also scored in line for focus and overexposure, and these readings update a running estimate, with its uncertainty, of the sample’s state or of where signal is likely to be found. A Bayesian belief map can also steer where to look next, with the choice made in code and no language model call; we refer to this as the “belief state”. No raw image or acquired data is passed to a language model.

#### Summarize

The cycle pass closes here. What the analysis returns is reduced to the few numbers the next pass needs, such as masks to counts, areas and intensities, each recorded with its source, so any value in the final answer traces back to the file and step that produced it. GENIE has a knowledge module which contains a static and extensible database of scientific knowledge, alongside self updating episodic memory and belief state. What goes to the knowledge modules is what the run established about how to run: which autonomy level suited a kind of goal, which failure is worth recalling, which exposure proved too high and more (Supplementary Note 3). The measurements themselves stay in the data. The summary returns to the planner, where the judgment to end or continue the cycle is made.

### Supplementary Note 2: On Language Model Calls

A language model is called where the work is interpretation: breaking down the goal, proposing what to do next, reading observations, writing analysis code, and judging whether a step is done. Between those calls the safety gate is fixed code, whose checks are enumerable. GENIE can use any language model, hosted by any provider or run locally; the live-cell experiment used Claude Opus 4.6 for the acquisition and Claude Fable 5.1 for the analysis, and the experiment on the second microscope used Claude Fable 5.1 only. This interpretive work is divided among nine agents, all served by the one model configured at the time (Fig. S1, Language model box). Each agent has its own instructions, its own inputs, its own generation settings and its own structured output, and no agent inherits another’s message history.

A communicator is bound to the user and runs only when the user sends a message, so it has no background thread and cannot reach the user on its own. A planner decomposes the goal into phases and each phase into subgoals, and runs in the mission thread. A deliberation engine runs continuously on its own thread, interpreting observations as they arrive and forming predictions about what should follow, so that an observation which violates a prediction is noticed. A code author writes analysis scripts, and a critic scores them against its own scientific validity rubric, which includes the statistical requirements of the prompt (replicate-level claims, effect sizes with uncertainty, stated assumptions), and audits their output for fabricated results. A router decides which knowledge modules a question belongs to; its calls are made with the least variability the model offers. An epistemic loop assesses candidate actions, discoveries, hypotheses and facts about the experiment and the system; its parts are a surprise detector, a hypothesis engine, a discovery ledger, a world model and a utility assessor.* A challenger, the least constrained of the agents, argues against the plan. A watcher agent, a further language model instance, follows the run from outside the mission thread, supplies hints to the planner and helps a generated script pass the critic; its messages enter the mission through the same channel as the user’s, become standing constraints, and are logged as such. Each agent’s calls carry a determinism level and a care level, which GENIE translates into what the provider offers: a sampling temperature where the model takes one (lowest for routing, highest for the challenger), or a reasoning-effort setting. For example in the mitochondria experiment, the acquisition model took temperatures; the analysis model took effort settings.

Several agents are instructed against inventing findings or confirmations, and prompting is not the only check. The deliberation engine may conclude that a field is empty rather than record a finding that is not there, and the communicator may not report an action as successful unless the function that ran it returned a result saying so. Beyond the prompts, generated code is audited against its own output for fabricated results, and the identities it claims to have written are checked against the data map. These reduce the chance of invention; they do not eliminate it.

Acquisition and analysis run on one set of machinery: a stage movement, a tile scan and a script GENIE writes are all named instructions drawn from the same permission table and logged the same way, differing only in which instrument or interpreter carries them out.

What GENIE knows is also held outside the model, in a body of knowledge it reaches through a fixed retrieval path. Most knowledge is static, several hundred domain modules spanning optics, fluorophore physics, phototoxicity and statistics, and anyone can read, correct or extend it. The rest accumulates as the system runs. What a mission establishes is written back as learned parameters, what went wrong is kept as an episode recalled when a similar situation arises, and anything anomalous is entered as a finding. A question may be answered by the modules alone, or a language model call may follow with the modules’ contents supplied to it. Which of the two happens is decided by what the modules return for that question and by a separate score of the answer’s quality: where they hold relevant content it is used as the answer, and where they hold little or none the model is called with whatever they did return. This body of knowledge holds the limits the gate checks against and the priors the planner starts from. It sits outside the language model’s context and can be inspected directly, and Supplementary Note 3 describes it in full.

*The epistemic loop is functional, and the behaviors described here are produced by the loop as a whole. Its surprise detector and world model are implemented and constructed at start up and are not yet fully realized; both are areas of active development.

### Supplementary Note 3: On the Knowledge Store

GENIE keeps its knowledge outside the language model’s context, in several separate stores (Fig. S1). Which stores a question is routed to is decided by the routing call and the lookup that follows is fixed code reading what those stores hold; where no store answers strongly, the language model supplies the answer instead. What a given step uses depends on what that step needs. The stores may answer it alone; the model may be called with their contents supplied to it; the model may be called without them where they hold nothing relevant; or neither may be consulted, as when an already validated script is run again. The effect is retrieval augmented, in that the stores supply grounding the model would otherwise have to produce on its own. The knowledge stores differ in where their contents come from. One is static, a textbook-like body of domain knowledge that ships with the system and that others can extend or correct; its modules were drafted with language model assistance and reviewed by the authors. A second is written as the system runs, holding what a mission established and what went wrong, kept as episodes that can be recalled when a similar situation arises, such as a grid scan that could not read the stage position on a previous run. A third records what the system finds, such as an accumulated exposure that is becoming damaging to a live sample. Accumulated exposure is the running total of illumination a sample has received over a mission, summed across every capture from the exposure time of each and held on the belief state, where the deliberation loop reads it; the per sample and per wavelength ceilings in the knowledge store are applied per capture rather than to the running total. Alongside these, a belief state holds GENIE’s running estimate, with its uncertainty, of things it cannot measure directly, for example the focus quality of the current field or which regions look worth imaging. Each such quantity is maintained as a Bayesian estimate, updated as observations arrive, and at higher autonomy levels these estimates feed a selection rule that chooses where to look next without the language model. The reported mission did not use this exploratory behavior (no L6 or L7 levels were engaged). A knowledge base holds fixed per sample settings the planner consults, such as an exposure ceiling or wavelengths flagged to avoid. A provenance record logs where every file came from and labels each raw file by the sample, replicate and channel it belongs to, so that later analysis can address data by what it is rather than by its filename.

Keeping knowledge in these stores rather than in the language model’s context, and admitting only numbers that trace back to a real measurement, reduce the chance of two of the ways a language model can fail over a long run at an instrument: losing track of its own state, and making things up.

### Supplementary Note 4: GENIE’s decisions

This note records the decisions GENIE made during the live-cell experiment, in the order of the supplementary figures, with the numbers from its own records. Supplementary Note 5 reports whether an independent computation agrees with each of them, and Supplementary Note 6 what the mission’s logs record of the run.

#### Acquisition and focus (Fig. S4a, b)

GENIE was told to autofocus every eight fields; its brightfield focus metric also triggered a refocus whenever it fell below threshold, 38, 20, 39 and 39 times in the four dishes, so it refocused more often than it had been told to, and the focal plane moved by 24, 57, 14 and 20 µm over a scan and was recovered at each event (Fig. S4a, b).

#### Illuminated field (Fig. S5a)

The illuminated field is a rectangle smaller than the sensor, off center, with a hard left edge and a soft roll-off toward the top, the bottom and the right, the same in every tile of every channel and dish. GENIE measured it from the mean profiles of all 2,296 tiles with a threshold of five standard deviations above the dead level and recorded rows 23 to 1199 and columns 205 to 1098 (Fig. S5a).

#### Erosion depth (Fig. S5b)

GENIE swept an erosion of the mask from 0 to 10 pixels (0 to 1.1 µm) and set a rule with two conditions, at least half of the mask pixels kept and a change of less than 2 % in the median field ratio, both to hold in every dish. Its erosion step kept 50, 45, 51 and 53 % of the mask pixels after one iteration, below the half its rule required in dish 2, so no depth from 0.11 µm (one pixel) on met the rule and GENIE operated at a depth of 0 (Fig. S5b).

#### The stage step

Fields of adjacent scan rows share content. GENIE measured the shift of that content by cross-correlation on 15 agreeing pairs, 978 pixels per recorded step of 89.76 µm, so the recorded stage positions are short by a factor of 1.1985, which it applied when placing the fields of a dish in one frame; the same correlation showed that image rows run along the stage’s Y axis. Column neighbors share no content, so GENIE took the stage’s own sign along X (Methods, “Placement”).

#### Distribution and classes (Figs. S7 and S9)

For each dish and each region size GENIE reported the histogram and the moments of the region ratio with a dip test of unimodality: every dish is unimodal at every size and right skewed (Fig. S7), with 2.6 µm region means of 3.07, 3.75, 4.12 and 3.79 (Methods, “Mitochondrial membrane potential distributions and classes”). GENIE defined HIGH and LOW regions as more than two standard deviations above or below the mean, which gave 369, 243, 714 and 2,031 HIGH and 0, 35, 54 and 109 LOW regions at 2.6 µm. The partner mean rises monotonically over the quintiles of the reference, the HIGH partners sit above random placement and the LOW partners below it (Fig. S9a, b), and the rings around low-ratio regions hold 10 to 15 % less mask than the rings around all regions, a density that does not order with the ratio the way the partner means do.

#### Region size (Fig. S8)

GENIE repeated the test at every region size (Fig. S8). The excess over the within-field null ends by 30 to 40 µm at every size (Fig. S8c), and the first-ring correlation and its excess are the same for regions from 0.66 to 21 µm (Fig. 2f).

#### Pre-exposure

Part of each window had already been lit once by the field of the previous scan row, and the prompt asked GENIE to check whether the ratio differs there. GENIE compared the two strips of the same field and read a small decrease, –0.006 to –0.13 log2 across dishes, below the 5 % change its own rule required to call a difference; the illumination roll-off across the field confounds that comparison, and the window rows, exposed once, are what every measurement uses.

#### The signal against acquisition time (Fig. S11a)

The prompt warned that beyond one field, distance in space is also distance in acquisition time. GENIE recorded the ratio of means of every field against its acquisition time (Fig. S11a) and reported a decline of ΔΨ_m_ over the scan. It considered the within-field null sufficient to account for this temporal effect and concluded that correlations remained distinguishable from the null at separations up to about 30 µm (Results).

### Supplementary Note 5: Independent verification of GENIE’s analysis

GENIE’s analysis was verified twice, by two authors who each worked from the raw tiles and used none of GENIE’s code. The expert verification (A) repeated GENIE’s image analysis from GENIE’s records of its settings and from its shared tables, with code written independently; where applicable, alternative approaches and estimators were tested against those GENIE used, to evaluate robustness. The recomputation (B) recomputed the mission’s outputs read-only from the raw tiles, with no file in the mission folder modified, and in addition measured what GENIE had left unmeasured. Each stands on its own. A is written as a methods section; B follows the order of Supplementary Note 4 and asks, for each decision GENIE made, whether an independent computation agrees with it.

#### A. Expert verification of GENIE’s image analysis

GENIE autonomously scripted an automated image analysis pipeline that extracted the relative fluorescence of two mitochondrial-associated fluorophores, mStayGold-COX8A and TMRE (Fig. 1; Methods, “Background, mask and erosion”). The pipeline then binned these pixels into regions of various sizes, valuing each region by its ratio of means, the mean TMRE intensity over its mask pixels divided by the mean COX8A intensity over the same pixels (Methods, “Regions”). These regional ratios were then correlated between reference points and their surroundings at various pixel distances in two sweeps, inside fields and across the whole dish (Fig. 2; Methods, “Spatial correlation and its nulls”). By using raw image files and consulting GENIE’s records, but not its scripted pipeline, we reproduced GENIE’s results for the within-field and whole-dish sweeps and reported similar coherence lengths, between ∼20 and 30 µm.

##### Within-field sweep

To verify GENIE’s within-field sweep, we first reproduced its segmentation and per-pixel ratio calculation. Consulting GENIE’s records, we adopted the same field exclusion criteria, the columns and rows of the illuminated sub-region to crop from the whole sensor image, and the masking algorithm and parameters. After reproducing the masks identically to GENIE’s analysis (about 32 million pixels across 967 fields), we calculated ratiometric pixel intensities for each masked pixel and compared them to those calculated by GENIE, confirming that any subsequent differences in analysis results were not due to differences in retained fields or masked pixels (Fig. S11b, c).

We then binned each image into non-overlapping square subregions with edge lengths of 24 pixels (2.6 µm) and calculated each region’s ratio of means from its background-corrected TMRE and COX8A intensities. As per GENIE’s records, we excluded regions comprising fewer than 25 masked pixels from subsequent analysis. For each retained region, we calculated the Pearson correlation coefficient between its own ratio of means and the ratio of means across a ring of regions surrounding the reference region. The ring of surrounding regions was maintained at a width of one region, but we calculated Pearson correlation coefficients between a reference region and rings at various distances from the reference region matching the distances swept by GENIE. For each replicate and separation distance, we compared reference-region and ring ratios in three ways:

Pooled correlation: a single Pearson coefficient calculated from all paired ratios across fields (Fig. S10a; Fig. S11c).

Field-centered pooled correlation: a single Pearson coefficient calculated across fields after subtracting a field’s mean reference-region ratio and mean ring ratio from their respective values. This metric verifies that the correlation trends persist after removing differences between fields (Fig. S11g).

Per-field correlations: a correlation computed within each field separately, summarized by the median and interquartile range (IQR) of these coefficients. This describes the typical within-field association and its variability across fields (Fig. S11c, d).

We report the pooled correlation, the field-centered correlations, and the median and IQR of the per-field correlations and found agreement with GENIE’s results (Fig. S10a; Fig. S11b–g).

Finally, we performed the null permutation analysis within each field to assess the floor that between-field variance imposes on the pooled Pearson correlation calculation. Following GENIE’s records, we shuffled the regions’ ratios of means either within or across fields, while each region’s position, masked-pixel count, and COX8A intensity remained in place to maintain ring membership, ring occupancy, and the total COX8A intensity of each ring. We performed permutations within each field 100 times per replicate and separation distance. We then repeated the within-field sweep on the permuted fields, and for each separation distance, we reported the null mean and standard deviation for the pooled correlation, the field-centered pooled correlation, and the median of the per-field correlations. Although our permutations differ from GENIE’s due to differences in the order in which we shuffled the ratios, we found strong agreement with GENIE’s reported null permutation test results (Fig. S10c).

##### Whole-dish sweep

To verify GENIE’s whole-dish sweep, we performed it in several ways. GENIE ran the whole-dish sweep across all the fields in a scanned replicate based on region tables containing exact frame positions. In addition to using GENIE’s approach to correlating regions and rings spanning multiple fields, we constructed a mosaic image of the scanned dishes by placing each field in its corresponding position on a lattice grid with 978-pixel spacing. In both approaches, we assumed 978-pixel spacing between fields, based on GENIE’s recorded measurement of overlapping content from adjacent fields. We replaced fields excluded from the within-field sweep with empty fields, but retained their positions within the compiled mosaic image. We then used the same approach employed in the within-field sweep, but swept the compiled mosaic images and extended the range of ring separations to reach about 1.35 mm, matching the separation distances GENIE tested. We reported the pooled Pearson correlation using the ratio of means of the TMRE and COX8A intensities over the rings’ retained regions. Plotting our results against GENIE’s showed strong alignment between the whole-dish sweeps, regardless of the differing approaches (Fig. S10f).

In addition to the whole-dish sweep GENIE performed, we also found the systematic decline in ratiometric pixel intensities that GENIE identified. By plotting the masked-pixel TMRE and COX8A intensities and their ratio over capture time (from imaging metadata), we reproduced GENIE’s finding (Fig. S11a). We then assessed the range of imaging periods that exhibited consistent negative declines by breaking the imaging sets into shorter intervals ranging from 1- to 30-minute blocks and computing Spearman correlations of each field’s ratio of means against time (Fig. S12a). We next calculated the percentage of intervals in which the ratiometric pixel intensity fell over time (Fig. S12b). We used these results to determine the maximum time difference between the first and last fields to include in subsequent whole-dish sweeps, choosing a conservative interval of 2.5 minutes between the first and last images of a multi-field sweep, which one row of the tiled imaging set that GENIE captured for each replicate (about 2.2 minutes) satisfies. We also performed this test across scanned areas broken roughly in half (about 0 to 20 min, rows 1 to 8, and 20 to 40 min, rows 10 to 17). We next ran the whole-dish sweep on these subsampled mosaic images as described above, and reported the same metrics (Fig. S12c). For the row-by-row whole-dish correlation, we observed an even steeper trend toward zero in the Pearson correlation coefficient starting at ∼10 µm, compared to the halved scanned areas. Finally, we repeated the same within-field null permutation test and reported the median across the subsampled mosaic images (Figs. S10e, S12d). The excess over the null decays at a distance of 32 µm regardless of approach or subsampled region size. Together, these analyses indicate that GENIE’s reported signal coherence lengths are robust.

#### B. Recomputation from the raw tiles Acquisition and focus (Fig. S4c)

In the recomputation, the fields flagged for defocus by the width of the COX8A structures, 2, 2, 2 and 0, hold under 1 % of the mask pixels, so no field needs to be excluded for focus (Fig. S4c).

##### Illuminated field (Fig. S5a)

The recomputation, with the same kind of rule held for 10 pixels, finds rows 29 to 1199 and columns 204 to 1098 (1171 by 895 pixels, 128.8 by 98.5 µm), within 6 pixels of GENIE’s rectangle, the difference at the top being the softer roll-off of the COX8A channel there (Fig. S5a).

##### Erosion depth and the mask rim (Fig. S5b, c)

Measured in one-pixel shells from the mask boundary, the ratio does not fall toward the boundary as a mixture with background would make it: the outermost shell sits within 7 % of the core, the second and third shells are 2 to 11 % above it, and the plateau starts at 0.44 µm (4 pixels) (Fig. S5b). The rim is a point-spread effect: at the same thin ridges the TMRE image is wider than the COX8A image by a median factor of 1.29 to 1.38 (65,075 ridges), in 99 to 100 % of fields, isotropic in both channels and with the stage at one read-back position for both exposures, and erosion cannot remove the rim without discarding every thin mitochondrion. GENIE’s erosion step eroded with a 3 by 3 square, which removes diagonal boundary pixels as well, so one iteration removes more than one pixel of depth; that is why one iteration kept only 50, 45, 51 and 53 % of the mask. Eroded by Euclidean distance, a depth of 0.11 µm (one pixel) keeps 55 to 62 % of the mask and removes the only shell where the floor acts on single pixels, and the recomputation operates there (Fig. S5b). The excess of the first-ring correlation over the within-field null for 2.6 µm regions is 0.30 at depth 0, 0.25 at 0.11 µm and 0.22 at 0.44 µm, above zero at every depth (paired t, p = 0.010 to 0.043): the conclusion does not depend on the erosion depth, the size of the excess does, by up to a third, and GENIE’s depth-0 figure is the upper end of that range (Fig. S5c).

##### The stage step

Fields of adjacent scan rows share content, and the shift of that content, measured on 169 pairs in the TMRE channel and 117 in the COX8A channel, is 982 pixels per recorded step of 89.76 µm (GENIE: 978 pixels from 15 pairs), 108 µm at 0.11 µm per pixel, where the record alone would give 816 pixels: the recorded stage positions are short by a factor of 1.203, and GENIE measured the same (1.1985) and applied it when placing the fields in one frame. The top rows of a field reappear at the bottom of the field in the next scan row, so image rows run along the stage’s Y axis, as GENIE found; the stage axes are aligned with the sensor within 0.3 degrees and the placement scatter of a field is about 8 pixels (0.9 µm). Column neighbors share no content, so the sign along X is an assumption, and running the whole-dish sweep under both signs changes no correlation by more than 0.05.

##### Distribution and classes (Fig. S9)

At GENIE’s settings the recomputation gives 2.6 µm region means of 2.99, 3.56, 4.07 and 3.78 (GENIE 3.07, 3.75, 4.12 and 3.79), unimodal in every dish and at every size and right skewed, with a long tail set by a small fraction of regions whose denominator sits just above its floor, which is why robust statistics are given next to the moments.Because the distribution is right skewed, the mean minus two standard deviations lies near zero and the LOW class is nearly empty by construction (0, 23, 65 and 122 regions against GENIE’s 0, 35, 54 and 109); with the median and a robust standard deviation both classes are populated.

##### Flat-field correction of the fluorescence ratio (Fig. S10b, h)

The ratio of the mean TMRE mask count to the mean COX8A mask count in 40-pixel blocks of the field, all fields of the four dishes pooled, runs from 0.80 to 1.17 between the 5th and the 95th percentile of blocks, the same in every dish and reproduced by the ratio of the two channels’ mean frame profiles: the instrument’s ratio is not flat across the frame. The flat-field correction divides every ratio by this pattern. It changes the first-ring excess by at most 0.02 and the coherence length not at all, lowers the region standard deviation by 3 to 6 %, and brings the share of the region variance that sits at fixed frame positions from 1 to 4 % to zero, against a field share of 21 to 60 % (Fig. S10b, h): the pattern is a small part of the field component and none of the result below one cell.

##### Region size and robustness below one cell (Fig. S10g, h)

In the recomputation the excess of the first-ring correlation over the within-field null holds at every region size from 0.66 to 10.6 µm (6 to 96 pixels), at erosion depths 0 to 0.44 µm, with and without the flat-field correction, with and without replicate 2 (N = 3, every sign holds and the p values roughly double), and under the pairwise estimator; at 2.6 µm it is 0.12 at 10 µm (p = 0.003) and 0.08 at 14 µm (p = 0.003) and gone from 18 to 25 µm. Holm’s correction across the rings of the sweep leaves the 10 and 14 µm rings significant (p = 0.02) with the first ring at p = 0.10, because the first ring’s excess varies most between dishes, 0.20 to 0.48 (Fig. S10g, h).

##### The region value (Fig. S13)

Figure 2 values every region and ring as the ratio of means over its masked pixels: the mean TMRE count over the masked pixels divided by the mean COX8A count over the same pixels. The recomputation reran GENIE’s whole-dish sweep, its two nulls and its class rule, with GENIE’s seeds and permutation rules, on GENIE’s own per-pixel ratio maps, changing only the value of a region to the mean of its pixel ratios (Fig. S13). The same code with GENIE’s value reproduces GENIE’s sweep and both nulls to 1.5 × 10^−8^, and with the mean of the pixel ratios the result is the same within rounding. The first-ring correlation is 0.71, 0.91, 0.91 and 0.79 with the ratio of means and 0.71, 0.90, 0.90 and 0.79 with the mean of the pixel ratios; the within-field null moves with it, so the excess is 0.44, 0.19, 0.34 and 0.21 against 0.44, 0.19, 0.32 and 0.21, with a mean of 0.29 under both (paired t, p = 0.015 and 0.015). The mean curve drops below the within-field null at 32 µm under both, and the HIGH and LOW classes keep their offsets from the dish mean at the first ring (3.88 and –3.44 against 4.06 and –3.65). The median of the pixel ratios gives the same result (excess 0.47, 0.19, 0.30 and 0.19, mean 0.29, p = 0.022).

##### Reproduction of GENIE’s numbers and what the verifications add (Figs. S10 to S12)

Recomputed with GENIE’s own method, GENIE’s numbers are reproduced. The expert verification, from GENIE’s shared tables, matches GENIE’s per-field ratio of means, mean and median pixel ratio, mask pixel counts and raw channel statistics exactly, and its sweep table to the file’s rounding (Fig. S11c); the pixel-ratio distributions agree within 0.007 at the quartiles (Fig. S11b) and the per-field ratios over time coincide (Fig. S11a). Its within-field sweep reproduces GENIE’s pooled correlation to the rounding of the shared file (Fig. S10a), the median over fields of each field’s own correlation (Fig. S11d), the first eight scan rows of the whole-dish sweep (median differences of 0.011 and 0.012 in the observed value and the null; Fig. S10e, S11e), the excess over the within-field null, which crosses zero at 24, 24, 29 and 28 µm in the verification and at 40, 23, 27 and 21 µm for GENIE (Fig. S11f), and the field-centered correlation of GENIE’s null step (Fig. S11g). The recomputation reproduces GENIE’s record at GENIE’s settings: the rectangle within 6 pixels, the durations, the fields with a mask within 3 %, the COX8A noise and floors, the median, mean and standard deviation of the region ratio at every size within 10 %, and the headline first-ring excess at 2.6 µm, 0.48, 0.19, 0.26 and 0.27 (mean 0.30, p = 0.017) against GENIE’s 0.44, 0.19, 0.34 and 0.21 (mean 0.29, p = 0.015) (Fig. S10a, g).

What the verifications add is what GENIE left standing. The expert verification reproduced the decline GENIE had reported: the ratio falls over every scan, by 1.34, 1.98, 1.49 and 1.91 % per minute, 41 to 55 % over a 35 to 40 min scan, while the COX8A signal stays flat, so the decline is in the potential-dependent channel alone (Fig. S12a). Cutting each scan into consecutive time blocks, it found that blocks of up to 2.5 to 3 min, about one grid row, hold no detectable trend (a change of at most 5 % in every dish at 2.5 min), and that from about 10 min the trend is resolved in every dish (Fig. S12b). As recorded, the whole-dish correlation between 2.6 µm regions and their ring stays above the whole-dish null to about a millimeter; on each half scan of eight rows it is lower, 0.30, 0.15 and 0.00 at 253 µm for the whole scan, rows 1 to 8 and rows 10 to 17 (Fig. S12c), and computed inside one grid row at a time, where every pair of fields was imaged within about 2.2 min, the long-range tail is gone and the excess over the within-field null, 0.51 at 2.6 µm, ends by 32 µm (Fig. S12d). Most of what GENIE reported as a field component reaching about 1 mm is therefore the decline of the potential-dependent signal over the scan, with a smaller part from the instrument pattern above and a residual at the scale of one or two fields that one scan per dish cannot separate from the dish. Acquisition time and grid row are collinear in these scans (Spearman rho 0.998), so a drift-free time block is also a contiguous band of the dish, and blocking by time does not by itself separate a temporal from a spatial cause; the half-scan comparison does, because the tail shrinks with the time span of the data at fixed geometry. The position pattern of the ratio and the erosion depth are the recomputation’s additions, above. Every conclusion GENIE drew below the scale of one cell stands. The standing limitation is N = 4 on every replicate-level test, at which the Wilcoxon floor is p = 0.125 and the first-ring excess rests on the t test. Software for the verifications: numpy, scipy, scikit-image, tifffile, matplotlib.

### Supplementary Note 6: Supplementary traces from the mission’s own logs (Fig. S3)

The guards described above leave a record when they fire, and this supplement reports what those records contain. Every count reported here and in Fig. S3 is a logged event. The sources are the five structured mission logs holding plans, subgoal rationales, gate verdicts and critic scores, the agent event streams and reasoning logs, the per output provenance records, and the 71 analysis scripts GENIE generated with their critique rounds. These were parsed by the scripts that built Fig. S3, which read the logs directly.

#### The run in full

The mission began with one natural language instruction on the microscope computer and acquired the four dishes in turn, each scan inside its 40-minute window (38.2, 35.6, 36.7 and 39.7 minutes from the first to the last exposure), 1,148 fields in all; the last eight fields of dish 2 were not acquired and the scan record was rebuilt from the tile headers, and GENIE waited between dishes for the operator to load the next. GENIE runs the analysis on the acquisition computer when nothing else needs the instrument; here the save directory was copied whole to a desktop with no instrument attached, to free the microscope for its next user, and the analysis continued there under the same goal from its own records. It opened with the image scale, the read noise and the data map in a two hour pass, continued after a settings update that raised the model’s token limits, and ran 28 hours of analysis over 37 cycles to its last accepted deliverable; it paused itself for 2.4 hours when the API credit ran out and resumed on its own, and that pause is not counted. That a mission can be relocated and continue where it left off is a feature of the design, not a requirement of it.

#### Convergence

Iteration is bounded and it converges. GENIE wrote 71 scripts over the analysis, of which 30 were accepted as steps and 36 were executed. Its code gauntlet logged 45 syntheses, 33 gate rejections against 26 regenerations that passed, 53 performance reviews and 71 fabrication audits that detected no fabrication, 129 critic assessments and 8 acceptances at the bar, and 31 rejected outputs quarantined rather than deleted. It refused to declare a phase complete 12 times, each time because the deliverables did not yet exist, and the cycle restarted until they did; plan failures numbered 12. The analysis made 1,374 calls to Claude Fable 5.1 and the acquisition 101 calls to Claude Opus 4.6, the two models configured for this mission; any hosted or local model can take their place. The rejections are substantive rather than cosmetic.

#### Provenance

Every accepted output carries its input MD5 and the script that made it, and every number in this manuscript’s results traces to one of those files; GENIE’s number index of 4,818 quantities, each with its record, is the source of the values quoted here. The text and the reference list of this manuscript were assembled by the authors.

### Supplementary Note 7: The full prompt

The complete instruction the operator gave GENIE at the start of the acquisition mission on the microscope computer, reproduced verbatim; the analysis continued under the same goal on a second computer (Supplementary Note 6; Fig. S3). The channel names are the instrument’s presets as the operator typed them: 561-S is the TMRE channel and 488-S the COX8A channel.

MISSION: Map mitochondrial membrane potential per unit mitochondrial mass across a 1.5 mm x 1.5 mm area of LIVE cells, then determine whether a region’s potential per unit mass predicts the potential per unit mass of other regions, beside it and far from it, and over what distances, from within a mitochondrion to a cell against its neighboring cells, across the full range of potential observed. 4 replicates (biological), loaded one at a time. Replicate 1 is already on the stage.

WHY TIME AND DOSE MATTER

The cells are live and the readout is potential-sensitive. Excess illumination bleaches the dyes AND depolarizes mitochondria: over-illumination does not merely degrade the measurement, it changes the quantity being measured. The dye is stable for a hard 40 minutes from the first exposure of each sample, and all acquisition for that replicate must finish inside that window. So: ACQUIRE FIRST. Start grid_scan the moment the mission starts. No pilot images, no parameter search, no test exposures, no rectangle detection, no analysis of any kind until all 4 replicates are acquired. Every acquisition setting below is given and final.

CHANNELS (given, do not re-derive)

- 561-S: the potential-DEPENDENT signal. 10 % power, 50 ms exposure.

- 488-S: the potential-INDEPENDENT mitochondrial mass signal (COX8A-labeled). 15 % power, 50 ms exposure.

- BF-Q: brightfield, for autofocus only. PHASE STRUCTURE (follow exactly)

acquire replicate 1 -> pause (operator loads replicate 2) -> acquire replicate 2 -> pause (operator loads replicate 3) -> acquire replicate 3 -> pause (operator loads replicate 4) -> acquire replicate 4 -> pause (operator copies the data off) -> analyze -> report.

Nothing from the analyze or report phases runs while any replicate is still to be acquired: analysis is compute-heavy, every sample is on a 40 minute clock, and nothing learned from an early replicate may reach the acquisition of a later one.

ACQUIRE (identical for every replicate) - replicate 1 is loaded, start acquisition right away.

1. Set laser powers (561-S 10 %, 488-S 15 %) and run ONE grid_scan call with: channels: [561-S, 488-S], exposures_ms: [50, 50], width_mm: 1.5, height_mm: 1.5, autofocus_every_tiles: 8, autofocus_channel: BF-Q, starting from the current stage position. That position is the recorded origin of the region.
2. Do not issue one grid_scan per channel, do not drop to fewer channels, and do not reorder channels for efficiency. At every position the order is 561-S, then 488-S, at every replicate, so the extra prior dose received by the second fluorescence channel is a constant bias, never a variable one.
3. grid_scan takes snapshots, which is what is needed. This is not a timelapse; nothing is measured over time. Do not take timelapses.
4. Record the stage position of every field, so that every measurement is locatable in one coordinate frame spanning the whole scanned region. Per-FOV pixel registration is done after acquisition, never during it.
6. Power and exposure are LOCKED. Apply the identical settings to all 4 replicates so values are comparable across them. Never re-optimize on replicates 2 to 4.
7. The area is fixed; time is the constraint. A 1.5 mm x 1.5 mm scan is expected to take about 40 minutes. If a replicate reaches 40 minutes from its first exposure with fields still unscanned, stop that replicate there and record which fields were not acquired. Do not shrink the area to save time, and do not extend beyond the window.

PAUSE (after each of replicates 1, 2, 3 and 4)

After replicates 1, 2 and 3 tell the operator: Replicate N acquired. Load replicate N+1 and confirm when it is on the stage. Wait for that confirmation. While waiting do nothing else: no analysis, no rectangle detection, no figures. After replicate 4 there is a pause too, and it is the last one: tell the operator: Replicate 4 acquired. All 4 replicates are acquired and the data can be copied off now. Confirm when the analysis may start. Wait for that confirmation however long it takes, and while waiting do nothing at all: no analysis, no rectangle detection, no figures, and no file of the acquisition is moved, renamed or rewritten. The analysis starts only on the operator’s confirmation, never on a timeout.

ANALYZE (only after all 4 replicates are acquired AND the operator has confirmed the pause that follows replicate 4; analysis may take as long as it needs)

A. IMAGE EXTENT, the first analysis step. The camera returns a 1200 x 1200 frame, but that is NOT the image. The illuminated field is a RECTANGLE smaller than the sensor, with hard edges rather than a gradual falloff, and it is not centered: the dead border on one side is wider than on the other. In recent acquisitions it spanned the full frame height and roughly 900 of the 1200 columns, with about 200 dead columns on one side and about 100 on the other, so roughly a quarter of every frame is dead. Do NOT assume those numbers. Measure the illuminated rectangle from the acquired images, per channel, report the edges found and how they were found, and exclude everything outside that rectangle from every step that touches pixels: background estimation, mask definition, the ratio, region definition and every comparison between regions. Including the dead border drags the background estimate down in both channels and places dead pixels among the neighbors of any region near the frame edge.
B. ACQUISITION QC FROM THE DATA ITSELF. Per replicate report: the fraction of saturated pixels in each channel, which are excluded from the ratio; how the signal changed over the scan, using each tile’s timestamp, and whether the 40 minute window held; the actual scan duration; the focus log; the fields reacquired or missing; and whether the channel exposed first at each position (561-S) is as sharp as the one exposed second, since a systematic deficit means the stage was still settling and that biases the ratio at every mask edge.
C. DEFINING THE MEASUREMENT. Membrane potential per unit mass is the ratio of the potential-dependent signal to the potential-independent signal. A ratio is only meaningful where both channels carry real signal from the same object, so three things are handled explicitly and each is justified from the data:

1. Background is removed from BOTH channels before any division. Estimate it independently in each channel from the image itself, never from a constant, and justify the level chosen from the measured noise distribution of that image.
2. The mitochondrial mask is defined from the potential-INDEPENDENT channel (488-S), because the region must not depend on the quantity being measured. Apply the identical mask to both channels.
3. Pixels near the EDGE OF THE MITOCHONDRIAL MASK are contaminated. This is about the mask boundary, not the edge of the image. Dim signal produces a less sharply defined and effectively more dilated mask, so pixels near that boundary hold a mixture of object and background in both channels. Their ratio is neither, and the mixture creates artificial gradients in the ratiometric image that can be mistaken for real structure. Erode the mask by a calibrated number of pixels and exclude what is removed. Determine that depth from the data rather than choosing it, by sweeping it: repeat the entire measurement across a range of depths, from none to well past what seems reasonable, and report every downstream result as a function of it. The sweep is not optional; it is also the check on erosion sensitivity that the statistics section requires. Once the depth is determined, apply that ONE value identically to every field and every replicate. The exclusion is a single consistent value across the whole sample, never adjusted per image or per field.

#### D. THE QUESTION

1. Build the map of membrane potential per unit mass across the whole scanned area, every measured region carrying its position in the single global coordinate frame. Do not stitch the fields into one image and do not register them to each other: a mosaic blends two exposures of the same sample wherever fields overlap, and nothing here needs one. Measure field by field, and place every region by the recorded stage position of its field. Pairs of regions that fit inside one field are formed inside that image, where their geometry is exact; only pairs that do not fit go through the stage positions, whose error is small against those distances; where both kinds of pair exist at the same separation, compare them. The fields overlap, because the scan steps by less than a field, and removing the dead border does not remove that: in the direction in which the illuminated rectangle spans the whole frame, about a third of every field is imaged again by its neighbor. The step is in the scan’s record, so nothing needs to be measured: give each field its own share of the grid, a window as wide as the step and inside the illuminated rectangle, the same window in every field, and count a region, a mitochondrion or a cell only in the field whose window holds its center, so that nothing is counted twice. The illuminated field beyond the window is not discarded: it is where the surroundings of a region near the window’s edge, and the rest of a mitochondrion or a cell that crosses it, are still seen in the same image. Whatever touches the edge of the illuminated rectangle is cut short: flag it, report how many, and keep cut cells out of every comparison that uses the cell as the unit. Part of each window had already been lit once by a field acquired before it: check whether the ratio differs there, and say so. Which image direction is stage +X and which is +Y, and their signs, is a property of the rig: settle it once from a few neighboring fields that share content, state it together with how far the recorded positions are off, and use it for every position. Report what fraction of the illuminated field ends up carrying a usable ratio, per replicate; if it is small, find out why before interpreting anything that follows.

2. DEFINE A REGION. Choose the region size, which MUST be larger than the optical resolution of the system, since anything smaller is not independently measured. State the size chosen, state the resolution it was checked against and how that resolution was determined, and repeat the analysis across a range of region sizes. At any one size, regions share no pixels. A region’s value is its potential per unit mass as a whole, the 561-S signal summed over its masked pixels divided by the 488-S signal summed over the same pixels, both after background removal, and not the mean of pixel-by-pixel ratios, which a dim denominator makes noisy and biased. Exclude regions that hold too few masked pixels, with the minimum justified from the data, and report how many were excluded.

3. THE DISTRIBUTION COMES FIRST. Do not reduce the map to categories before showing what is there. Bin region potential per unit mass across its full observed range and report the resulting distribution: the histogram, the mean, the standard deviation, the shape, and whether it is unimodal. Report this per replicate, and pooled for display. Everything below is computed across the FULL range of bins, not only at the extremes.

4. THE CLASSES ARE A SLICE OF THE DISTRIBUTION, NOT A REPLACEMENT FOR IT. From that distribution define HIGH as more than 2 standard deviations above the mean and LOW as more than 2 standard deviations below it. Report both thresholds and how many regions fall in each. Then repeat across a range of thresholds and report every result as a function of threshold, so no conclusion rests on 2 sigma alone. High and low are reported as the sharpest cases of the general result, never instead of it.

5. THE TEST. Determine whether the potential per unit mass of a region tells you anything about the potential per unit mass of OTHER regions, beside it and far from it, across the full binned range and additionally for the HIGH and LOW classes specifically. What is compared is always potential per unit mass, never morphology or raw intensity. The sweep is over SEPARATION: pair each region with the regions found at a given distance from it, collected in rings of a stated inner and outer radius in micrometers, counting only what lies inside the eroded mask, and sweep that distance without gaps, several rings to each doubling of the distance, from just beyond the optical resolution up to the size of the whole scanned area, using the global coordinate frame to pair regions across fields. Do not stop the sweep at the scale of a mitochondrion: it is meant to walk through the scales of the biology, from within a mitochondrion, to neighboring mitochondria, to a whole cell, to a cell against its neighboring cells. Use rings, not growing windows: a window that always contains the nearest neighbors cannot say at what distance a relationship lives. Do not define the separation by distance from a mitochondrion: mitochondrial shape is variable, sometimes punctate and sometimes tubular or networked, and a distance from a mitochondrion is ambiguous because it depends on whether it is measured from the mask edge, the centerline or the center of mass. A ring centered on the region has none of that ambiguity. Distance alone does not say which structure two regions share, and the mask does. A mitochondrion is a connected piece of the mask. A CELL can be traced from its mitochondria alone: a group of masks gathered around a void, the nucleus, and ending where the mitochondria end, the way an eye picks out a cell in the 488-S image. Define cells that way, from the 488-S mask and at a scale determined from the data, show the traced cells over the image so they can be judged by eye, and use the voids as the check: a traced cell holding several voids, or none, is probably not one cell, so report how often that happens. Then carry the comparison up the levels: at matched separations compare pairs on the same mitochondrion, on different mitochondria of the same cell, and in different cells; repeat the test with the mitochondrion and then the cell as the unit, a cell against its neighboring cells; and report how the total variation in potential per unit mass divides among regions within a mitochondrion, mitochondria within a cell, cells within a field, and fields. Express every relationship as how much the other regions change per unit change of the region, a slope with its uncertainty, and not only as a correlation coefficient: a coefficient against an average also depends on how many regions went into that average, which changes with distance and with how much mask a ring holds. Report how much mask each ring holds, and where too little is left report the measurement as uninformative rather than reporting a value.

6. THE NULL. Test the observed relationship against regions drawn at random from the same map, so that any structure reported is structure the null does not produce. Report the outcome with uncertainty. If regions are no more predictive of other regions than random regions are, at any bin, report that plainly. The null keeps the mask, the positions and the rings exactly as they are and reassigns only the values, at every level, region size and separation, and the observed result is always shown next to it: what the geometry alone produces is not zero, and it must be visible. Reassign the values once within each field and once across the whole replicate: the difference between the two is what the fields’ own differences contribute, and it is not to be read as the sample’s.

7. SEPARATION. Report every result as a function of the separation distance between the paired regions, and include separations small enough to fall within a single mitochondrion. Do not exclude or discount short separations: mitochondrial membrane potential is reported to be heterogeneous at all scales, including within a single mitochondrion, so a within-mitochondrion result is a real result and not an artifact of proximity. Report the short-separation and long-separation regimes distinctly rather than pooling across them, and let the data say where, if anywhere, the relationship changes. For each bin, and for HIGH and LOW, also show the mean potential per unit mass of the other regions against separation, relative to the replicate’s mean; the departure from that mean is the effect size. As a control, show the same way how much mask the rings around each class hold: if HIGH and LOW regions sit in differently dense surroundings, say so, because density then competes with potential as the explanation. For each replicate, report the separation beyond which the relationship can no longer be told apart from the null, and the one beyond which HIGH and LOW no longer differ: one distance per replicate, on which the replicate-level claim about reach rests. Beyond one field, distance in space is also distance in acquisition time, and every field carries the instrument’s own pattern: separate both from the sample before attributing anything at those scales to it.

#### E. STATISTICS

The biological replicate is the unit of inference: N = 4, with per-replicate n = the regions measured. Compute per-region measurements, aggregate to a per-replicate summary, and make every claim at the replicate level. Do NOT run inferential tests on pooled regions.

- Every parameter entering the measurement must be determined by something measured in this run, not chosen, inherited or assumed. For each, report the measurement that determined it, the range examined, and the result at both ends of that range. If a parameter cannot be determined from the data, do not pick a plausible value: report it as undetermined and say what would be needed.

- Where a test is repeated across separations, bins, region sizes or any other swept quantity, account for the number of tests.

- Report effect sizes and uncertainty, not only p-values. State the assumptions behind any test and justify parametric or non-parametric.

- Where the design or N = 4 limits what can be claimed, say so plainly.

REPORT

Overview potential map: for each replicate, ONE map of the potential per unit mass over the whole scanned area, at the erosion depth applied to the measurement. It is for the eye and for the figures; no measurement is taken from it. Assemble it from the per-field ratio maps by placing each field’s window at its recorded stage position, in the orientation settled in D.1, with no blending and no image-based refinement of the positions, and leave it empty where there is no data. Write per replicate: the map at full resolution as a float32 TIFF if it fits in memory as a single array, and if it does not, say so; the map binned 4 x 4 (mean of the valid pixels of each block) as a float32 TIFF; a placement record listing every field with the position used and whether it was included or excluded and why; and a PNG at 600 dpi or more on ONE color scale common to the four replicates (pooled 1st to 99th percentile of the ratio), gray where there is no data, with an appropriate scale bar, a round length in micrometers suited to the extent shown, labeled with that length, and computed from the pixel size in the acquisition record and from whatever binning or resampling the displayed map went through, never from an assumed value. State the placement uncertainty in the caption.

Plots, SuperPlot convention: per-region data colored by replicate, per-replicate summaries overlaid as larger points; any inferential test runs on replicate-level values. Curves follow the same convention: each replicate as its own thin line, the mean across replicates with its uncertainty as a thick one. Show the four overview potential maps side by side on their common color scale, show the binned distribution of potential per unit mass, and show every swept quantity as a curve over the swept parameter, never as a single chosen value: erosion depth, region size, threshold and separation distance. Give every size, radius and separation in micrometers as well as pixels, never in units of another swept parameter. Show every result that runs over separation on a logarithmic distance axis, and again in separate panels for each range of scales (within a mitochondrion, across a cell, between cells, across the scanned area), each with its own vertical range: a trend at one scale is invisible on an axis set by another. Annotate every plot with test name, statistic, p-value, effect size and N at the level the test was run. Save the numbers behind every curve and every figure, per replicate, in a file next to it, not only counts or summaries. Write every figure at 600 dpi or more, and give every image a scale bar computed the same way as the overview map’s.

Text and figures, publication-ready: answer every question above completely, report every mask, every intermediate, report every swept quantity as a full curve, and do not omit a question because its answer is negative or inconclusive. If a question cannot be answered from the data acquired, say so and say what would be needed. Enumerate every statistical test: assumptions, test chosen and why, statistic, p-value, effect size, N. Report the acquisition settings used and the time budget actually held for each replicate. For every acquisition and analysis decision, state the method and justify it from data. Flag any output that looks unreliable rather than reporting it as final. Log every failure and the decision taken to solve or circumvent it. This should be a ready to submit to a peer-reviewed scientific journal.

---

**Figure S1.**
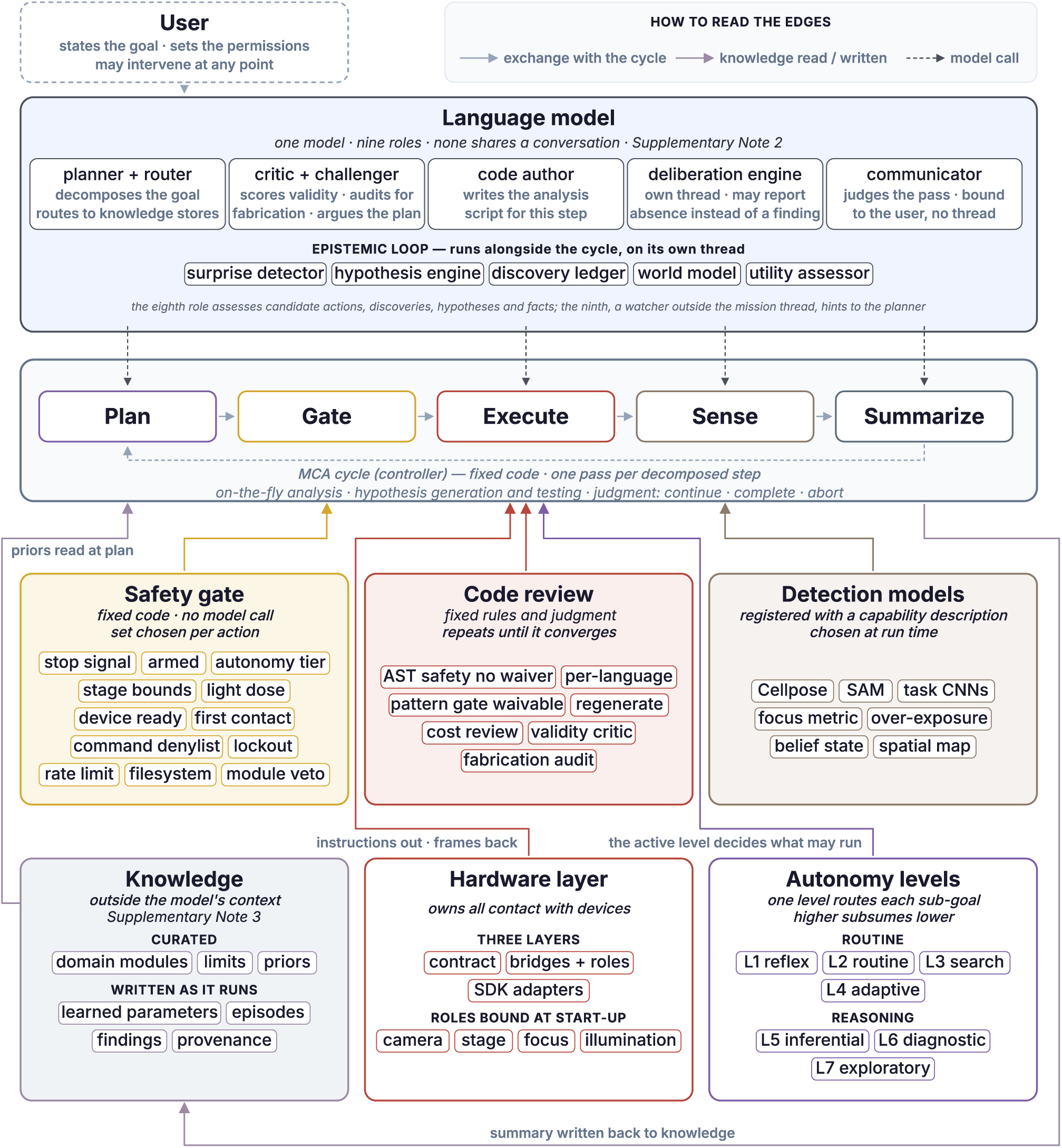
Each stage of GENIE’s cycle draws on its own models, checks and stores. An expanded view of the system sketched in Figure 1, in the colors of Figure 1a and b. At the top is the language model in its nine roles (Supplementary Note 2): planner and router, critic and challenger, code author, deliberation engine, communicator, a watcher that follows the run from outside the mission thread, and the epistemic loop, which runs on its own thread and assesses candidate actions, discoveries, hypotheses and facts with a surprise detector, a hypothesis engine, a discovery ledger, a world model and a utility assessor. The watcher agent, a further language model instance, supplies hints and helps a generated script pass the critic through the same channels as the user. In the center is the controller’s five-stage cycle, fixed code that runs one pass per step of the plan and closes with continue, complete or abort. Below it are the safety gate (deterministic checks assembled per action, no model call), code review (the screens a generated script passes until it converges), the detection models (perception models registered by capability), and the three subsystems the cycle consults: the knowledge stores, curated and run-written; the hardware layer, which owns all contact with devices; and the seven autonomy levels. Arrows connect each box to the stage it serves: priors are read at plan, the summary is written back to knowledge, the hardware layer exchanges instructions and frames with execute, and the active autonomy level determines what may run.

**Figure S2.**
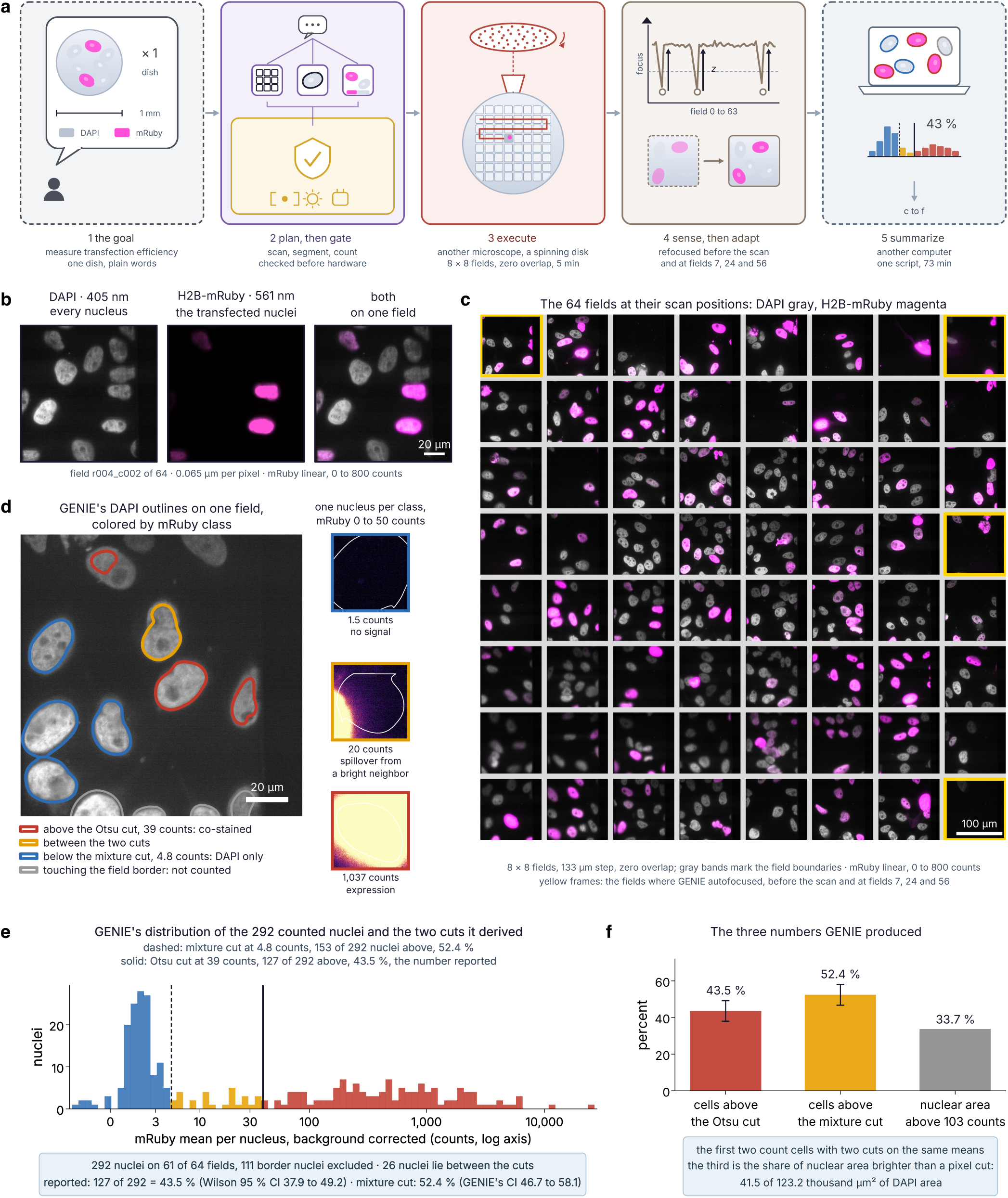
GENIE runs autonomously on a second microscope with entirely different hardware and answers a new question. (a) The run on GENIE’s cycle. The goal was the transfection efficiency of H2B-mRuby in one dish of fixed, DAPI-stained cells; GENIE planned behind the safety gate, drove a spinning disk microscope through its hardware layer, scanned 8 by 8 fields at a 133 µm step with no overlap in 5.2 min, refocusing before the scan and at fields 7, 24 and 56, and ran the analysis on another computer with one script in 73 min (schematic drawn with AI assistance). (b) One field three ways: every nucleus (DAPI, 405 nm), the transfected nuclei (H2B-mRuby, 561 nm), and both. (c) The 64 fields at their scan positions, DAPI in gray and H2B-mRuby in magenta; gray bands mark the field boundaries and yellow frames the fields where GENIE autofocused. (d) GENIE’s DAPI outlines on one field, colored by the mRuby class of each nucleus against GENIE’s two cuts, with one nucleus of each class on one mRuby scale at right: no signal, spillover from a bright neighboring nucleus, and expression. (e) GENIE’s distribution of the 292 nuclei it counted on 61 of 64 fields (111 border nuclei excluded) and the two cuts it derived: 127 nuclei lie above the Otsu cut, the transfection efficiency reported (43.5 %), and 153 above the mixture cut (52.4 %). (f) The three numbers GENIE produced; the third is the share of nuclear area above a pixel cut, 33.7 %. Scale bars, 20 µm (b, d) and 100 µm (c). Pixel size, 0.065 µm.

**Figure S3.**
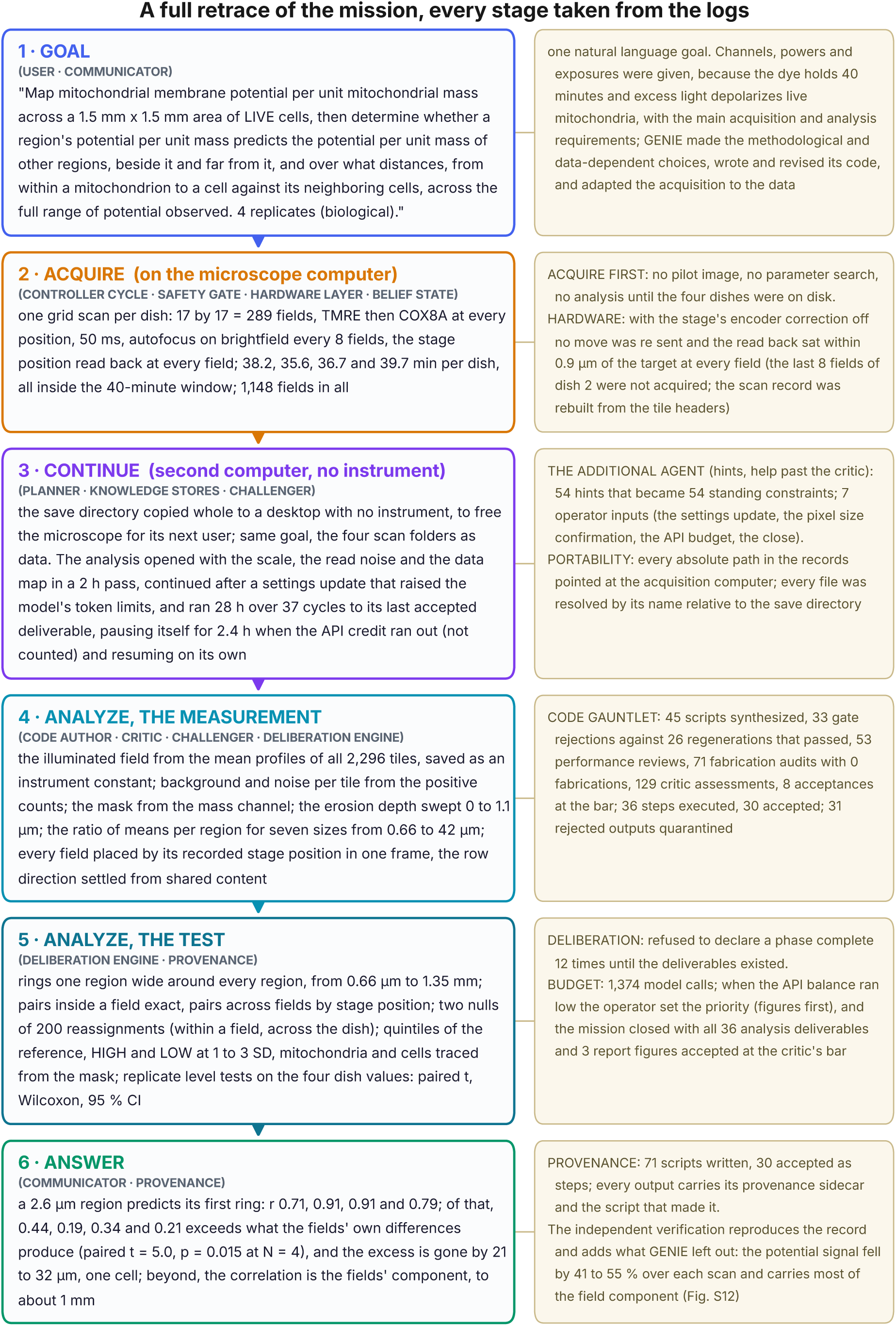
The mission’s logs retrace it stage by stage. The run reconstructed from the mission’s own records in six stages, each annotated with the guards that fired and how often, counted from the mission logs. 1, Goal. One natural language instruction named the sample, channels, powers and exposures and the main acquisition and analysis requirements; GENIE made the methodological and data-dependent choices, wrote and revised its code, and adapted the acquisition to the data (the complete instruction is reproduced in Supplementary Note 7). 2, Acquire, on the microscope computer: one 17 by 17 grid scan per dish, the TMRE channel then the COX8A channel at every field, autofocus on brightfield every eight fields and the stage position read back at every field; 38.2, 35.6, 36.7 and 39.7 min, 1,148 fields in all. The last eight fields of dish 2 were not acquired and the scan record was rebuilt from the tile headers. 3, Continue the analysis. The save directory was copied to a desktop with no instrument, and the analysis continued there under the same goal from its own records: a 2 h opening pass, then 28 h over 37 cycles to its last accepted deliverable, with a 2.4 h pause when the API credit ran out. The watcher agent supplied 54 hints that became standing constraints, and the operator gave 7 inputs (a settings update, the pixel size confirmation, the API budget, the close). 4, Analyze, the measurement: the illuminated field from all 2,296 tiles, the background and noise per tile, the mask from the mass channel, the erosion sweep, the ratio of means for seven region sizes and the place of every field in one frame; 45 scripts synthesized, 33 gate rejections against 26 regenerations that passed, 71 fabrication audits with none detected, 129 critic assessments, 36 steps executed and 30 accepted, 31 rejected outputs quarantined. 5, Analyze, the test: rings one region wide from 0.66 µm to 1.35 mm, two nulls of 200 reassignments, quintiles and the HIGH and LOW classes, mitochondria and cells traced from the mask, and replicate-level tests on the four dish values; 12 refusals to declare a phase complete before its deliverables existed, 1,374 model calls, and all 36 analysis deliverables and 3 report figures accepted at the critic’s bar. 6, Answer. A 2.6 µm region predicts its first ring (r 0.71, 0.91, 0.91 and 0.79, of which 0.44, 0.19, 0.34 and 0.21 exceeds the within-field null; paired t = 5.0, p = 0.015), the excess is gone by 21 to 32 µm, about one cell diameter, and beyond that the correlation is the fields’ component. The verifications (Supplementary Note 5; Figs. S4c, S5 and S10 to S13) reproduce the record and add the decline with acquisition time, 41 to 55 % over each scan, which carries most of the field component.

**Figure S4.**
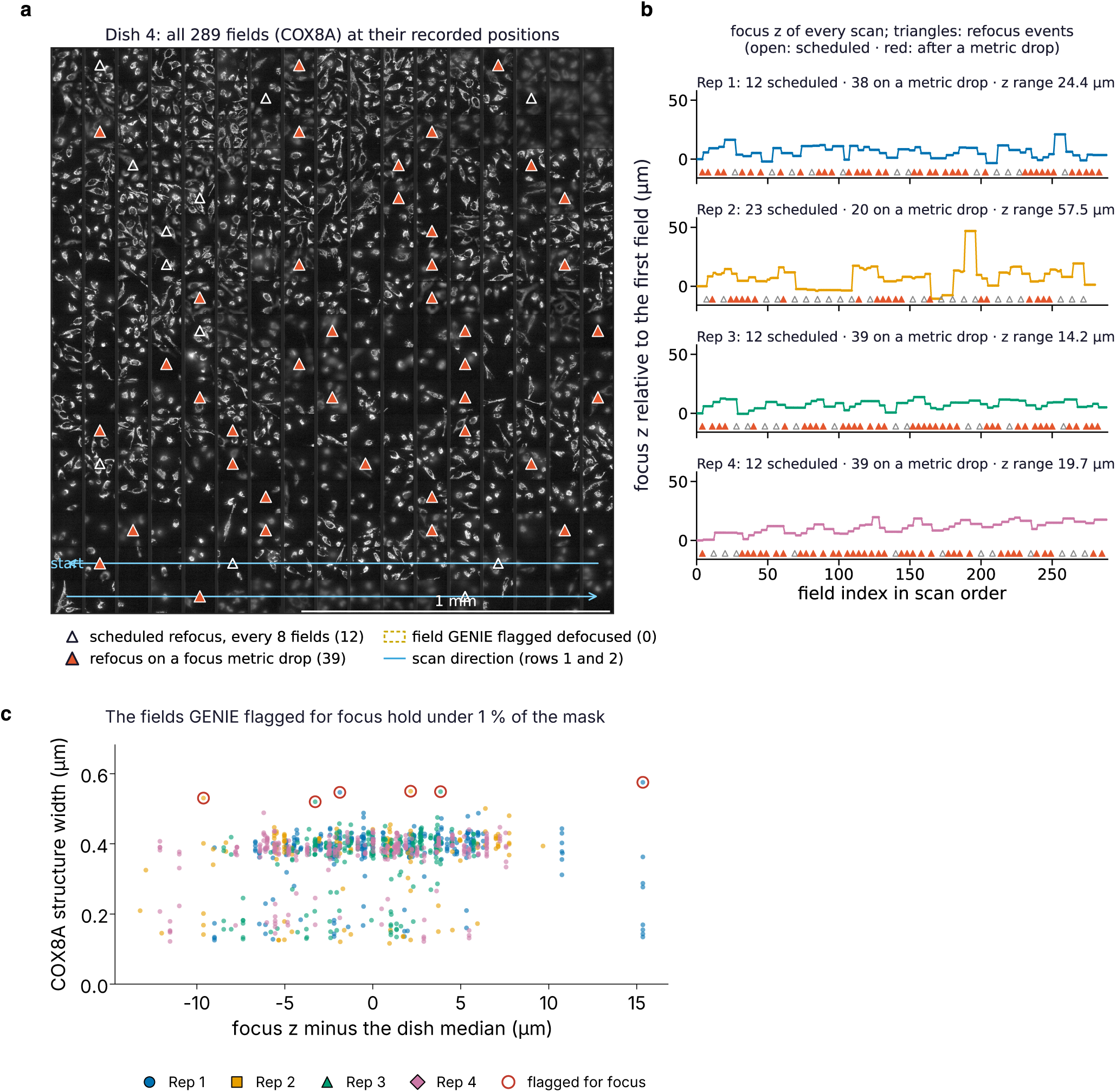
The scans held focus, as a full run at its recorded positions and every refocus show. (a) The COX8A image of all 289 fields of dish 4 placed at GENIE’s recorded stage positions (its own canvas, 0.88 µm per displayed pixel), with the scan direction of the first two rows (blue) and every refocus GENIE ran, marked on the field after which it ran: open triangles the scheduled refocus every eight fields (12), red triangles a refocus triggered because the brightfield focus metric fell below its threshold (39). Dashed yellow would mark fields GENIE flagged as defocused; this dish has none. The dark stripes between columns are the sample the stage step leaves unimaged. Scale bar, 1 mm. (b) The focus z of every scan relative to its first field against the field index in scan order, one panel per dish, with the same triangles; the counts are 12 scheduled and 38 on a metric drop, 23 and 20, 12 and 39, and 12 and 39, and the z ranges are 24.4, 57.5, 14.2 and 19.7 µm. Most refocusing was triggered by the metric, and the scans held focus within the depth of the cells throughout. (c) From the recomputation (Supplementary Note 5), the width of the COX8A structures in each field against that field’s focus z minus the dish median, one color per dish; the fields GENIE flagged for focus (red rings) hold under 1 % of the mask. The field profiles and the timing of the scans are in Figure 1c.

**Figure S5.**
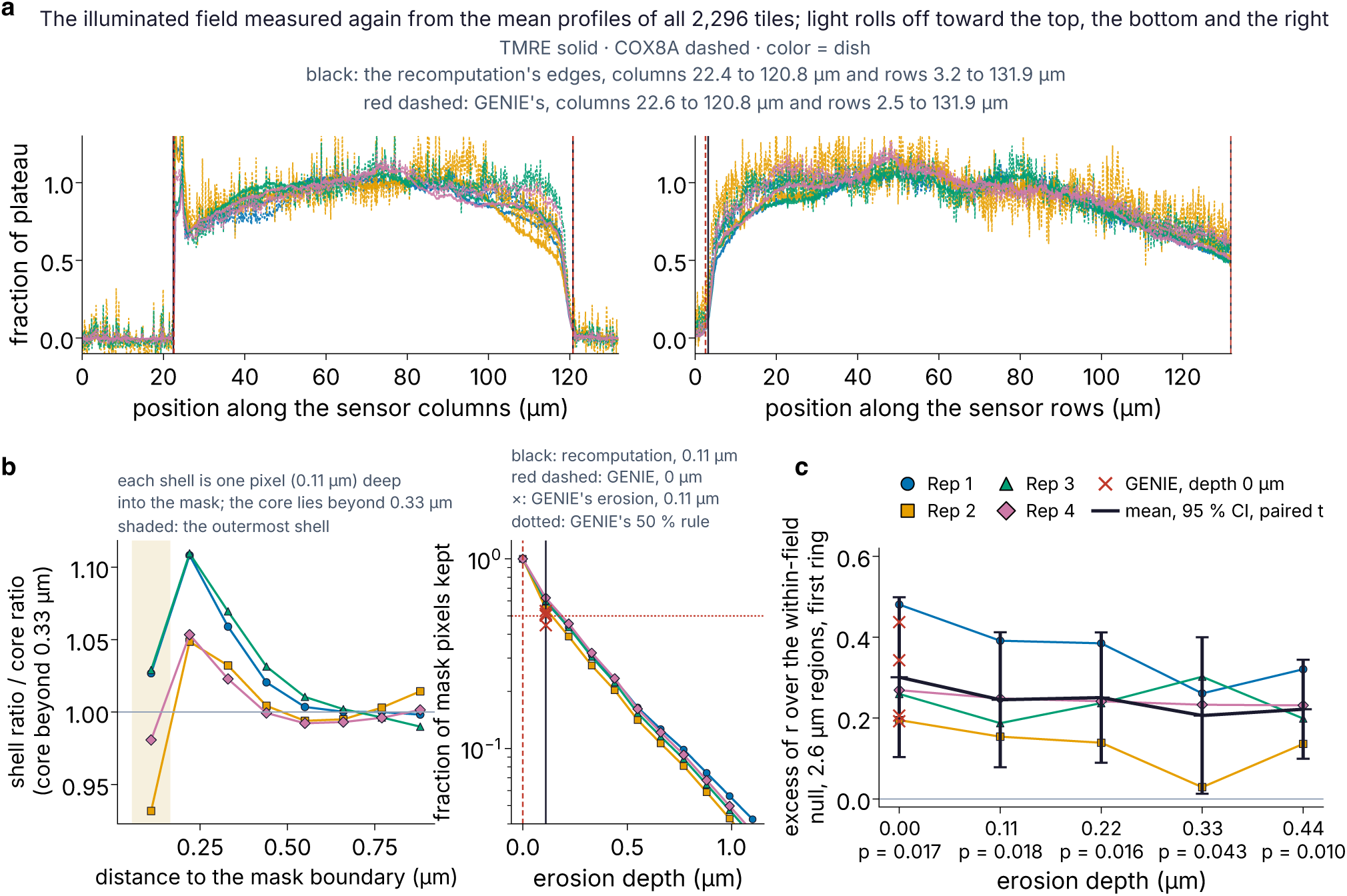
The illuminated field and the erosion depth: what GENIE determined, checked from the raw tiles. Every curve in this figure was computed by the recomputation from the raw tiles (Supplementary Note 5), with code written for the check and none of GENIE’s; GENIE’s own recorded values are drawn in red for comparison. (a) The mean profile over every tile of each dish in each channel, along the columns (left) and the rows (right), as a fraction of the plateau (TMRE solid, COX8A dashed, one color per dish). The recomputation sets an edge where the profile leaves the dead level by five standard deviations of the dead border: columns 22.4 to 120.8 µm and rows 3.2 to 131.9 µm of the sensor (black). GENIE’s record (red dashed) agrees within 0.7 µm. (b) Left, a measurement GENIE did not make: the mask cut into shells one pixel (0.11 µm) wide, counted inward from its boundary, each point the ratio of means of one shell divided by that of the core deeper than 0.33 µm, per dish. The outermost shell (shaded; 38 to 45 % of the mask) sits at 0.93 to 1.03 of the core and the next two at 1.02 to 1.11; from 0.44 µm inward the shells sit at the core’s value. The rim is a few percent high because the TMRE image is softer than the COX8A image, and erosion cannot remove it without discarding every thin mitochondrion. Right, the fraction of mask pixels kept against erosion depth, per dish, for erosion by Euclidean distance as the recomputation does it (colored lines); the recomputation operates at 0.11 µm (black line), which keeps 55 to 62 % of the mask. GENIE’s erosion removes one pixel from every side at each iteration (a 3 by 3 square), which cuts more: one iteration kept 50, 45, 51 and 53 % (red crosses), below its own rule of one half (red dotted) in dish 2, so GENIE operated at depth 0 (red dashed). (c) The excess of the first-ring correlation over the within-field null for 2.6 µm regions, recomputed at each depth, per dish, with the mean over the four dishes and its 95 % confidence interval (paired t, p under each depth), and GENIE’s values at depth 0 overlaid (red crosses; 0.44, 0.19, 0.34 and 0.21). The mean excess is 0.30 at depth 0, 0.25 at 0.11 and 0.22 µm, 0.21 at 0.33 µm and 0.22 at 0.44 µm, above zero at every depth (p = 0.010 to 0.043).

**Figure S6.**
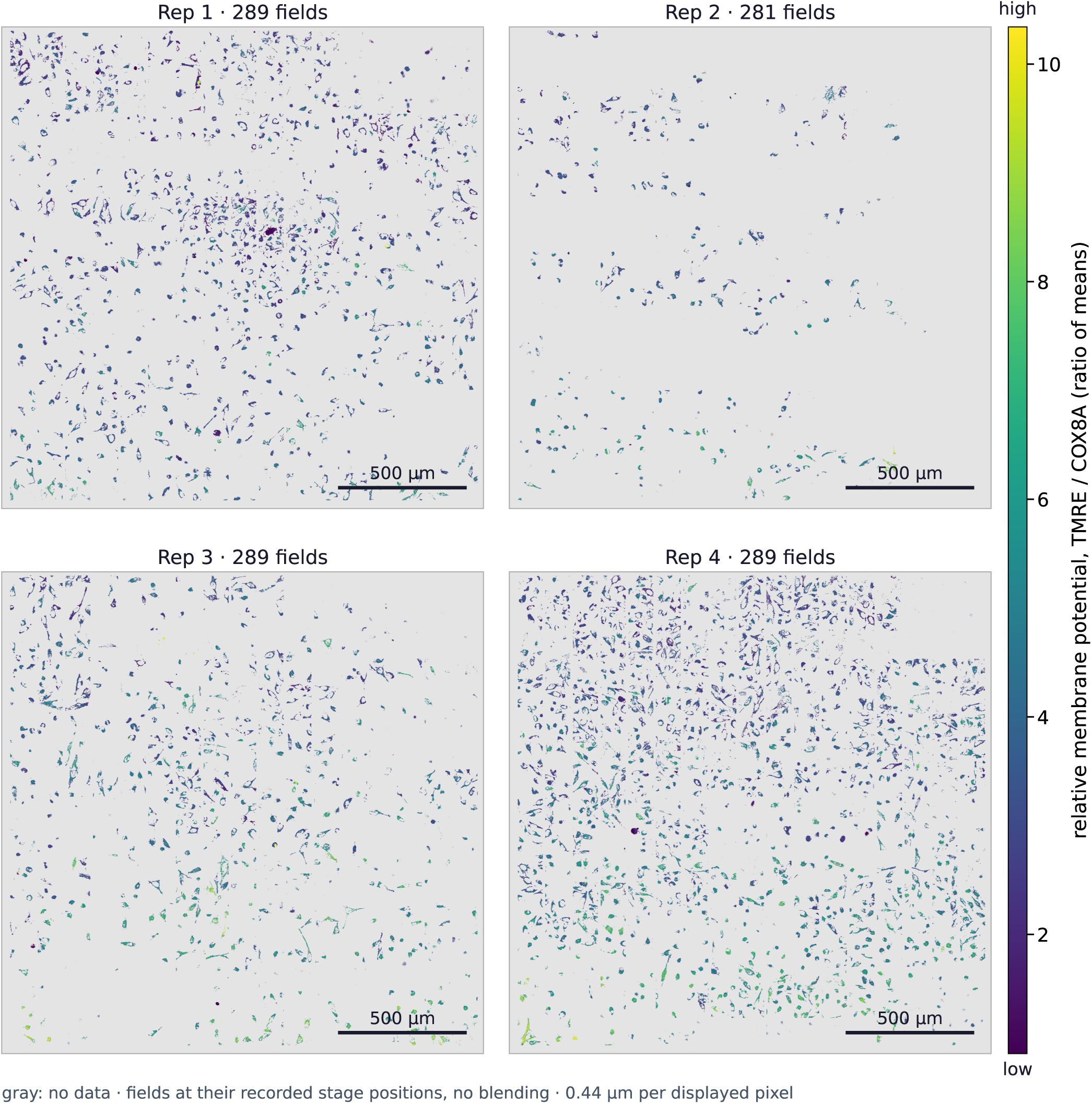
GENIE mapped relative mitochondrial membrane potential for the four dishes on one color scale. GENIE’s overview maps for replicates 1 to 4 (289, 281, 289 and 289 fields): its 4 by 4 binned canvases at 0.44 µm per displayed pixel, each field at its recorded stage position with no blending, and gray where no pixel carried a ratio. The color runs from low (purple, 0.91) to high (yellow, 10.3), GENIE’s pooled 1st to 99th percentile over the four dishes. The first scan row is at the bottom of each map, and the eight fields of dish 2 that were not acquired leave its empty corner at the top. Figure 1d shows replicate 4 alone. Scale bars, 500 µm.

**Figure S7.**
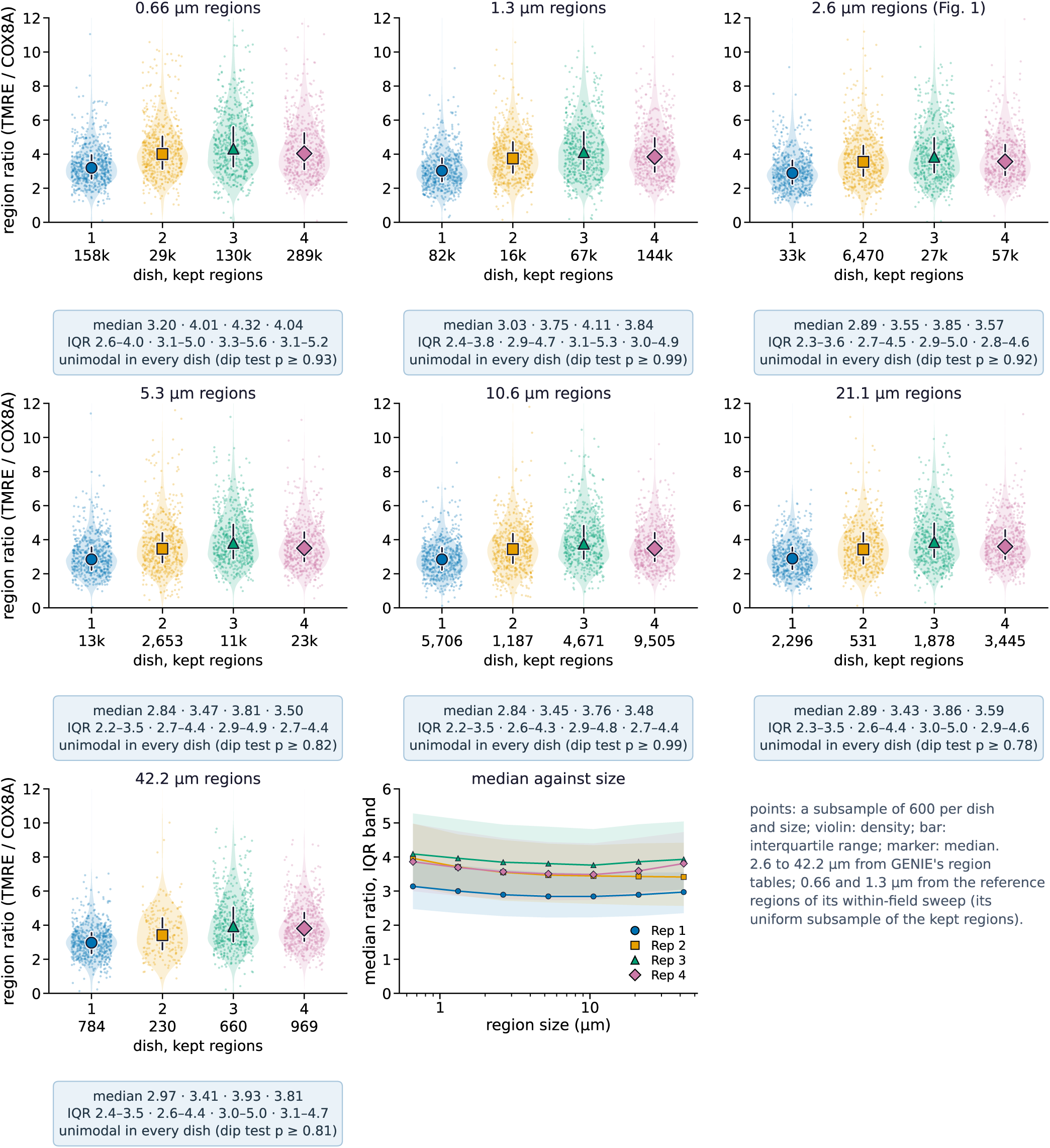
GENIE’s region ratio is unimodal in every dish at every region size it swept. SuperPlots of the ratio of means (TMRE over COX8A) of GENIE’s kept regions of the retained fields, one panel per region size from 0.66 to 42 µm: a subsample of 600 points per dish, the violin as the density, the bar as the interquartile range, the marker as the median, and each dish’s number of kept regions under it (GENIE’s distribution summary). Every dish is unimodal at every size (Hartigan dip test p ≥ 0.78) and right skewed, and the medians move by under 15 % across a 64-fold change in region size (last panel, the median against size with the interquartile band). Values at 2.6 to 42 µm are from GENIE’s region tables; at 0.66 and 1.3 µm, where it wrote no region table, from the reference regions of its within-field sweep. The pixel distribution is in Figure 1f.

**Figure S8.**
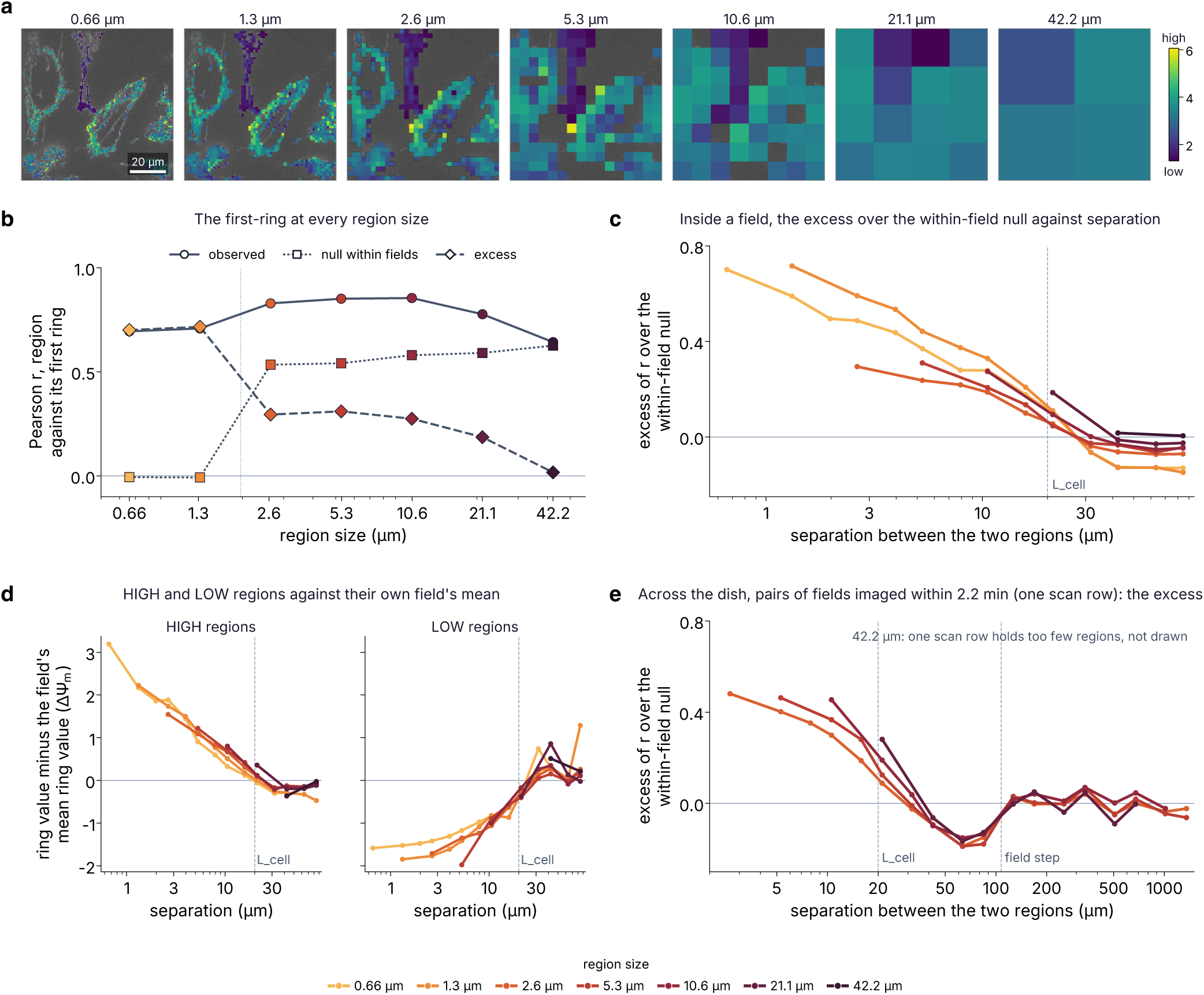
GENIE ran the same test at every region size it swept, from 0.66 to 42.2 µm. Each curve is the mean of the four dishes at one region size, colored from the smallest size (light) to the largest (dark); the per-dish values are in the numbers file. (a) GENIE’s regions on one field of replicate 4 (field 282) at the seven sizes, on the same 84.5 µm square of the field, colored from low (1.3) to high (6.1). (b) The first ring against region size: the observed correlation between a region and its first ring, the null that reshuffles values within each field, and the excess of the first over the second. From 2.6 µm on, the excess is 0.29, 0.31, 0.27 and 0.19 and falls to 0.02 at 42.2 µm, where the first ring already lies 42 µm away. Below 2.6 µm GENIE ran only its within-field sweep, whose correlation removes every field’s mean first (dotted line), so its null sits at zero and the excess equals the observed value (0.70 and 0.72). (c) The excess over the within-field null against separation inside a field, from GENIE’s records; it is gone by 32 µm at every size up to 5.3 µm and by 42 µm at 10.6 and 21.1 µm. (d) The ring partners of HIGH and LOW regions minus the mean of their own field, as in Figure 2d: at every size up to 10.6 µm HIGH regions start above their field and LOW regions below it, and both reach their field’s mean by 21 to 32 µm. (e) The excess over the within-field null across the dish for pairs of fields imaged within 2.2 min, one scan row at a time, as in Figure 2e: GENIE’s estimator and its within-field null on every grid row, the median over rows per dish, and the mean of the dishes. At the first ring the excess is 0.48, 0.46, 0.45 and 0.28 for 2.6, 5.3, 10.6 and 21.1 µm regions, and it is gone by 32 µm (2.6 and 5.3 µm) and 42 µm (10.6 and 21.1 µm). A ring resting on fewer than 100 reference regions in a row is left out, and at 42.2 µm one scan row holds too few regions, so that size is not drawn. Guides mark L_cell, GENIE’s median area-equivalent cell diameter (20 µm), and the field step (107.6 µm). Panels d and e were computed by us from GENIE’s regions with GENIE’s rules.

**Figure S9.**
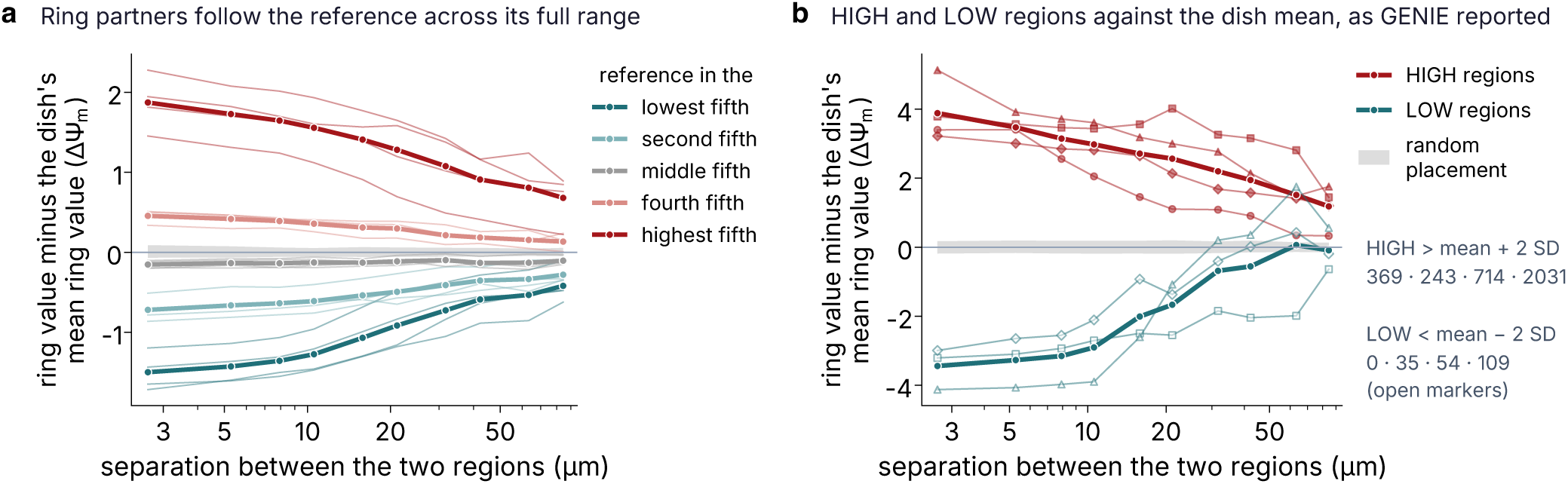
The ring partners across the range of the reference, as GENIE reported them. GENIE’s records for 2.6 µm regions, with the ring partners compared with the dish mean. (a) The mean value of the ring partners minus the replicate mean against separation, for reference regions in each fifth of the dish distribution. (b) The same for HIGH regions, more than two standard deviations above the mean (369, 243, 714 and 2,031 regions), and LOW regions, more than two below (0, 35, 54 and 109; dish 1 has none, so it has no LOW curve), against the band of random placement. Thick lines are the mean of the four dishes and thin lines the dishes. Figure 2d shows the same classes against the mean of their own field.

**Figure S10.**
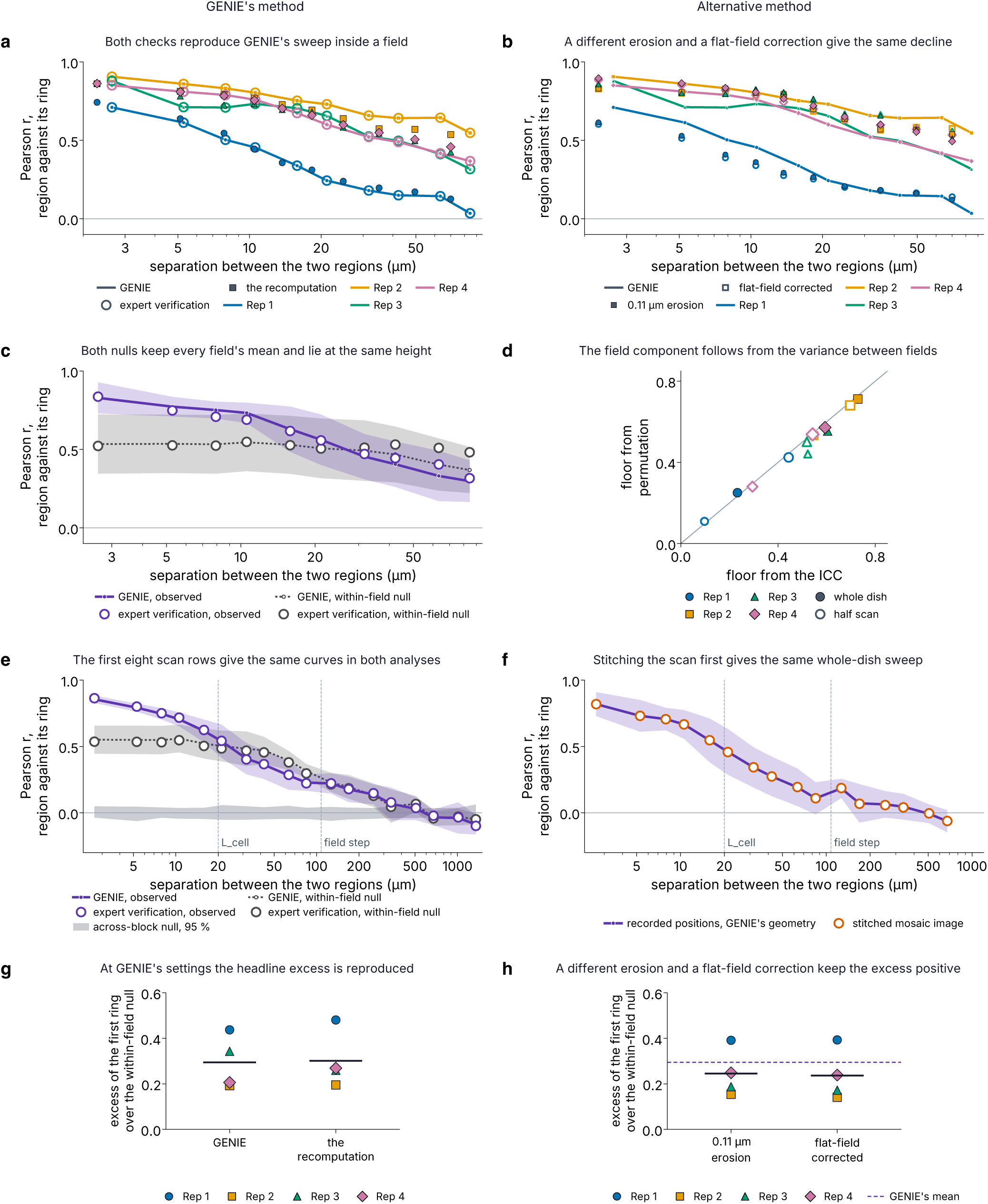
GENIE’s method and an alternative method reach the same result for every quantity where both were run. GENIE’s method (left column) repeats what GENIE did, step by step, to check the math; an alternative method (right column) measures the same quantity another way. The checks come from the expert verification, written independently of GENIE and plotted from its shared tables, and from the recomputation from the raw tiles (Supplementary Note 5). Every panel uses 2.6 µm regions; lines are GENIE’s records unless stated. (a) The sweep inside a field: GENIE’s pooled correlation between a region and its ring (lines, one color per dish), the expert verification’s (open circles), which matches it to the rounding of the shared file, and the recomputation at GENIE’s settings (small filled markers), which follows it within a median of 0.04 (largest 0.11; its rings are centered slightly differently). (b) The same sweep in the recomputation with 0.11 µm erosion (filled) and with the flat-field correction (open): the first ring moves from 0.71, 0.91, 0.88 and 0.85 to 0.61, 0.83, 0.86 and 0.90 and to 0.60, 0.83, 0.86 and 0.89, and the decline with separation keeps its shape. (c) The within-field null: observed correlation and null as the mean of the four dishes with bands of ± 1 SD, GENIE’s (lines) and the expert verification’s (open circles); at 2.6 µm the null is 0.53 for GENIE and 0.52 for the verification. (d) The height of that null predicted from the variance between fields alone (an intraclass correlation, no permutation) against the height the permutation gives, for the four dishes (filled) and eight half scans (open); the median gap is 0.017 and the largest 0.081. (e) The whole-dish sweep on the first eight scan rows, GENIE’s estimator on GENIE’s recorded field positions (lines, mean ± 1 SD across dishes, with the null reshuffled across the block in gray) and the expert verification on the same regions and geometry (open circles); over 72 values the observed correlation differs by a median of 0.011 and the within-field null by 0.012. (f) The same sweep with the fields first stitched into one mosaic image, a step GENIE never took, against the sweep on GENIE’s recorded positions (line), as the mean over the eight half scans with a band of ± 1 SD; the two differ by a median of 0.002 and at most 0.029. Guides in e and f mark L_cell, GENIE’s median area-equivalent cell diameter (about 20 µm), and the field step. (g) The headline excess of the first ring over the within-field null for 2.6 µm regions, from GENIE’s record (0.44, 0.19, 0.34 and 0.21; mean 0.29) and from the recomputation at GENIE’s settings (0.48, 0.19, 0.26 and 0.27; mean 0.30). (h) The same excess with 0.11 µm erosion (0.39, 0.15, 0.19 and 0.25; mean 0.25) and with the flat-field correction (0.39, 0.14, 0.17 and 0.24; mean 0.24), against GENIE’s mean (dashed); the excess stays positive in every dish.

**Figure S11.**
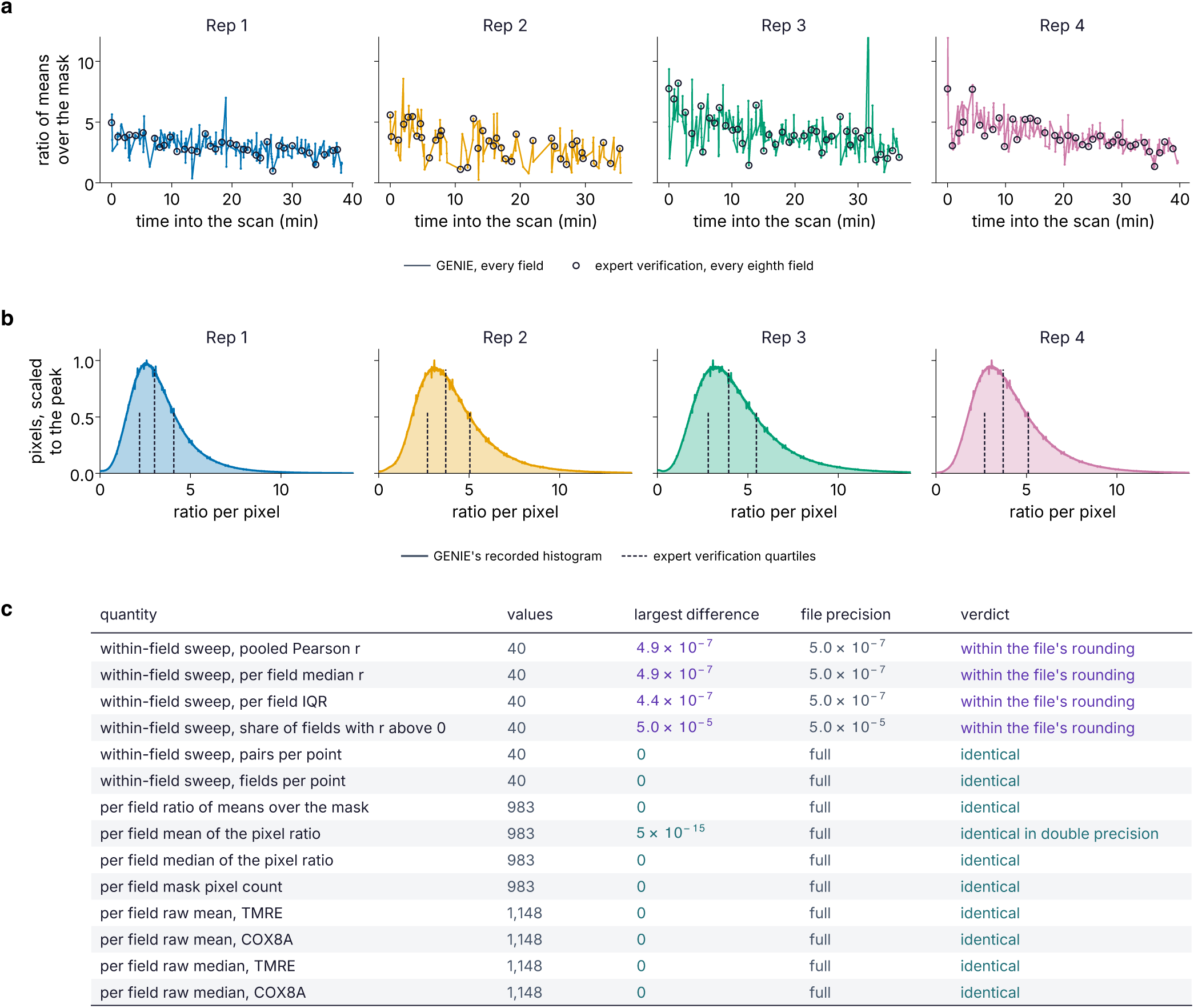
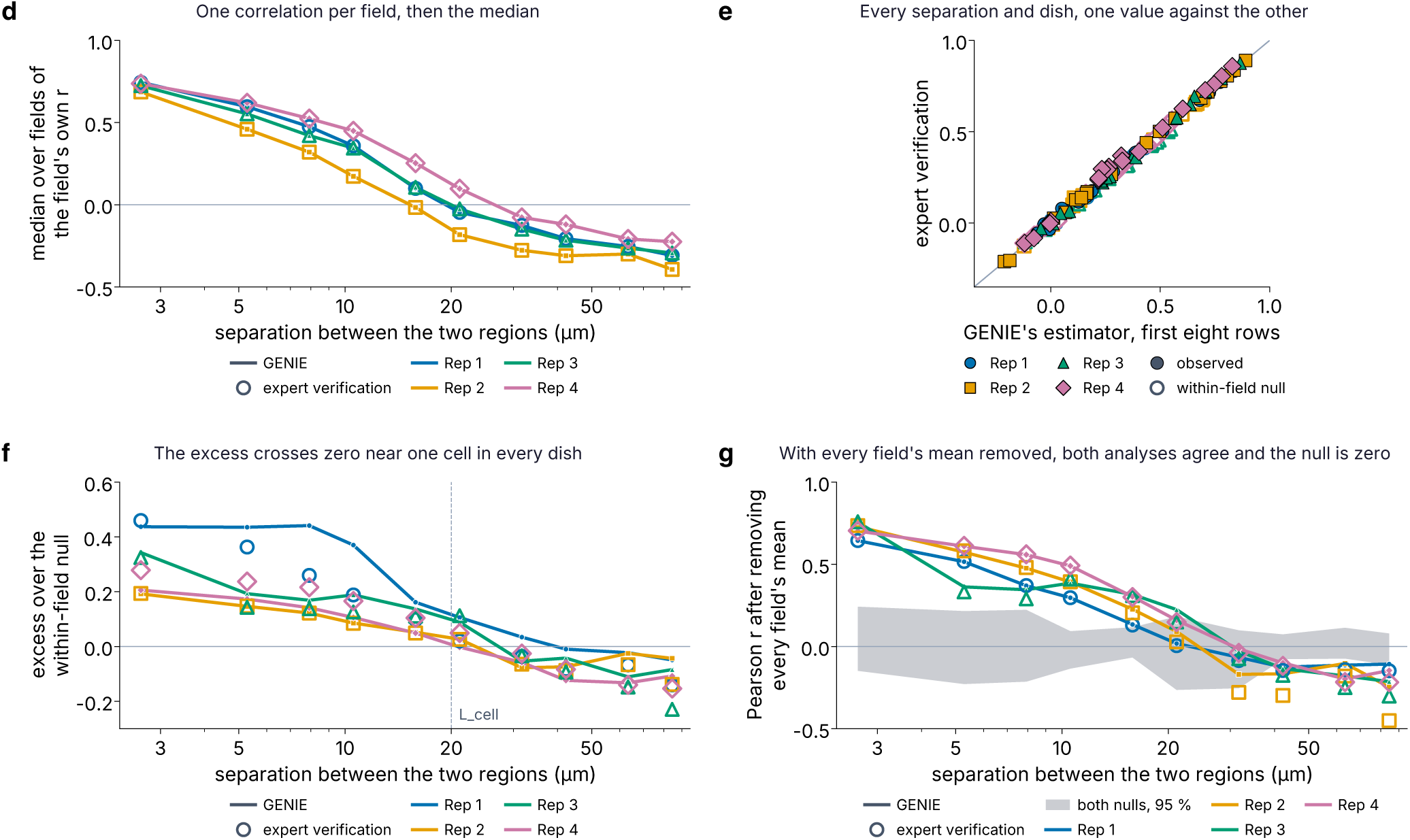
Recomputed with GENIE’s own method by the expert verification, GENIE’s numbers are reproduced. Every panel repeats GENIE’s own method to check the math, from the expert verification’s shared tables. (a) The ratio of means over the mitochondrial mask for every field against the time into the scan, GENIE’s (lines) and the expert verification’s (open circles, every eighth field shown). (b) GENIE’s recorded pixel-by-pixel ratio distribution per dish with the expert verification’s quartiles (dashed), which agree within 0.007. (c) Every quantity the expert verification compared, 8,764 values in all, with the largest difference from GENIE’s value and the precision to which the shared file was written: the tables written at full precision agree exactly, and the correlation table agrees to its sixth decimal. (d) GENIE’s median over fields of each field’s own correlation (lines) and the expert verification’s (open markers), 2.6 µm regions; they agree to the file’s rounding. (e) The first eight scan rows of the whole-dish sweep, every dish and separation, the expert verification’s value against the sweep of Figure S10e, observed (filled) and within-field null (open); median differences 0.011 and 0.012. (f) The excess of the correlation over the within-field null, per dish, GENIE’s (lines) and the expert verification’s (open markers), with L_cell dashed; the excess crosses zero at 24, 24, 29 and 28 µm in the verification and at 40, 23, 27 and 21 µm for GENIE, whose sweep also pairs regions across fields. (g) The statistic of GENIE’s own null step, the correlation after every field’s mean is removed, GENIE’s (lines) and the expert verification’s (open markers), with both nulls (gray, 95 %); up to 10.6 µm they agree within 0.008 in dishes 1, 2 and 4 and within 0.055 in dish 3.

**Figure S12.**
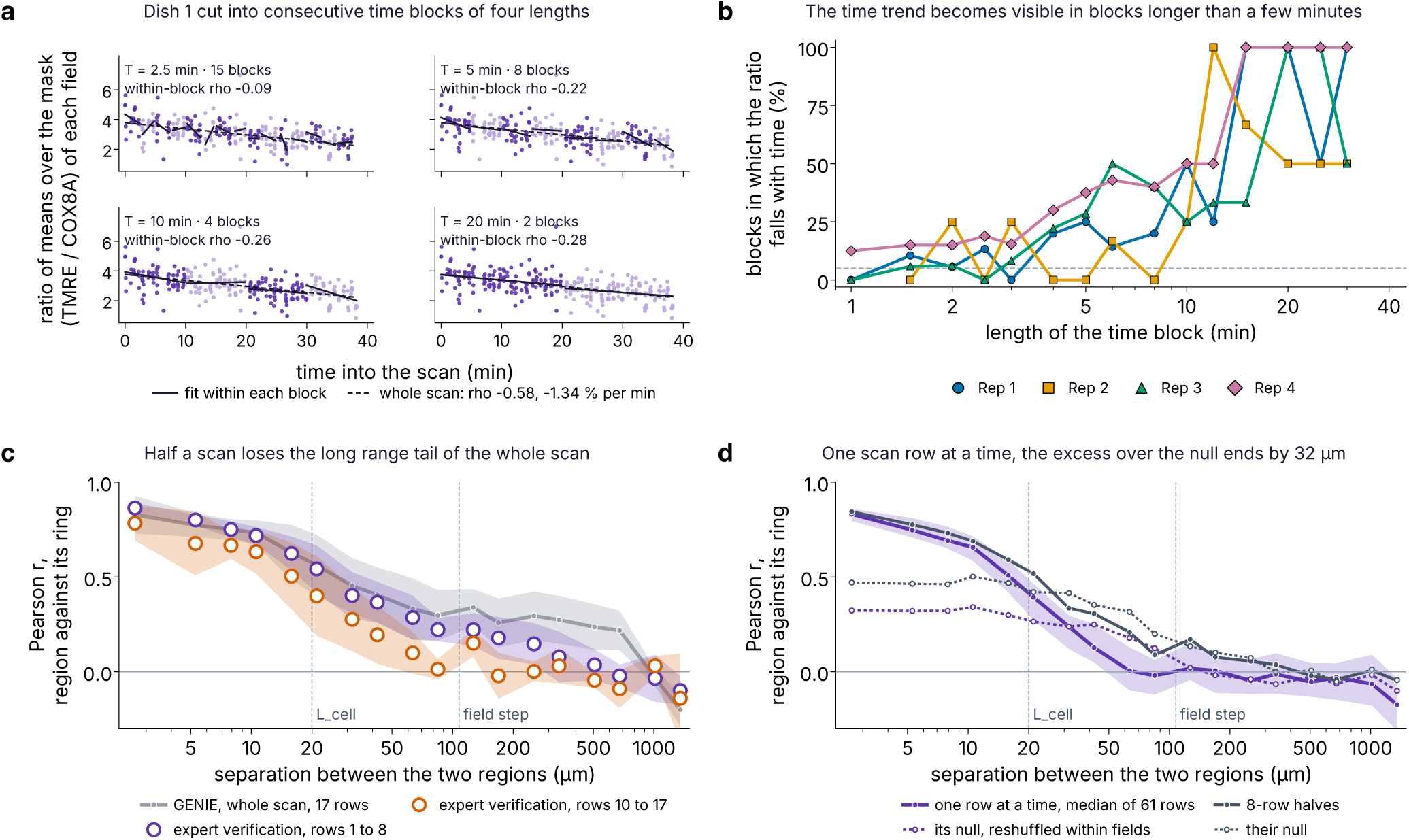
The potential-dependent signal against acquisition time, by the expert verification. Every panel is from the expert verification’s tables, drawn here. (a) The ratio of means over the mask of every field of dish 1 (277 fields over 38.2 min) against the time into the scan, cut into consecutive time blocks of 2.5, 5, 10 and 20 min as the expert verification cut it (a block needs at least 10 fields). Points alternate in tone from one block to the next, each block carries its own least-squares line, and the whole-scan fit is dashed (Spearman rho −0.58; −1.34 % of the mean ratio per minute). Each panel gives the median within-block rho and the block count: −0.09 over 15 blocks of 2.5 min, −0.22 over 8 blocks of 5 min, −0.26 over 4 blocks of 10 min and −0.28 over 2 blocks of 20 min. The expert verification used its own mask here; its whole-scan slopes are −1.34, −1.98, −1.49 and −1.91 % per minute, a fall of 41, 53, 43 and 55 % over the four scans. (b) The share of blocks in which the ratio falls significantly with time (p < 0.05) against the block length, one line per dish, with 5 % dashed: blocks of up to 2.5 to 3 min, about one grid row, show no consistent trend, and from about 10 min the trend is resolved in every dish. (c) The whole-dish sweep on each half scan of eight rows against GENIE’s whole scan of 17 rows, mean ± 1 SD across dishes: at 253 µm the whole scan gives 0.30, rows 1 to 8 give 0.15 and rows 10 to 17 give 0.00; guides mark L_cell and the field step. (d) The whole-dish sweep computed inside one grid row at a time, where every pair of fields was imaged within about 2.2 min, so the time trend is removed by construction rather than corrected: the median of the 61 rows (purple, band the interquartile range) and its null reshuffled within fields (dotted), against the 8-row halves and their null (gray). The null drops by 0.08 to 0.18 at every separation, and the excess over it, 0.51 at 2.6 µm, is gone by 32 µm (29 µm with GENIE’s median estimator). The same construction with GENIE’s own estimator and nulls is Figure 2e.

**Figure S13.**
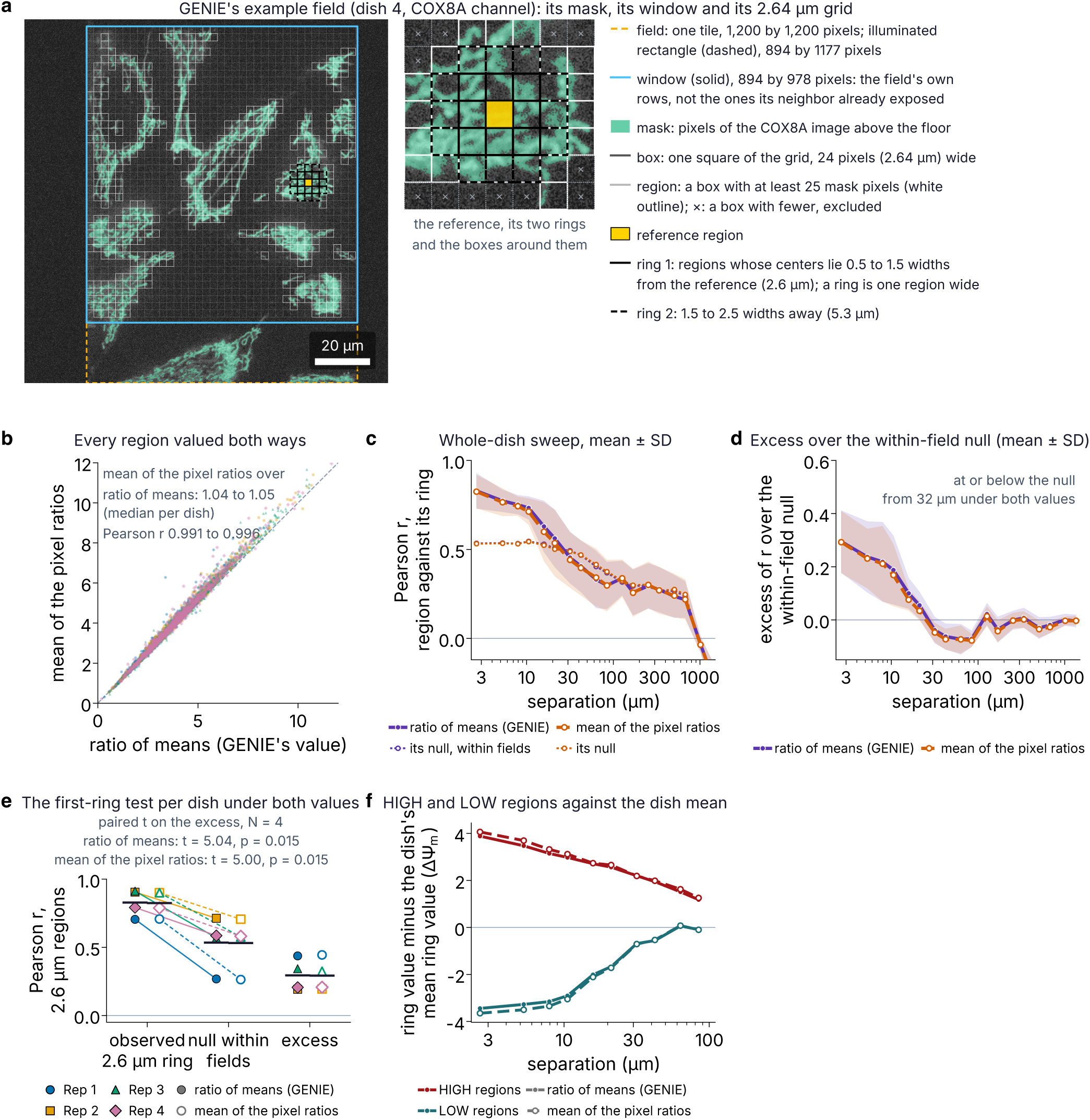
The region value: GENIE’s ratio of means against the mean of the pixel ratios. (a) The vocabulary of the analysis on GENIE’s example field (dish 4, COX8A channel). The field is one camera tile, 1,200 by 1,200 pixels; the lasers illuminate a rectangle of 894 by 1,177 pixels (dashed), and the window (solid) is the 894 by 978 pixels of it that the field owns, the rows its neighbor in the scan does not expose again. The mask (green) is the set of pixels of the COX8A image above the floor. The grid cuts the window into boxes of 24 by 24 pixels (2.64 µm); a box with at least 25 mask pixels is a region (white outline) and a box with fewer is excluded (×); this field holds 537 regions. Right, the reference region (yellow) with its first ring (solid) and its second ring (dashed), the regions whose centers lie 0.5 to 1.5 and 1.5 to 2.5 box widths away. Scale bar, 20 µm. (b) Every region valued both ways, 1,500 regions per dish drawn: GENIE’s ratio of means (the mean TMRE count over the region’s mask pixels divided by the mean COX8A count over the same pixels) against the mean of its pixel ratios. The second is larger by a factor of 1.04 to 1.05 at the median, and the two correlate at r = 0.991 to 0.996 per dish. (c to f) GENIE’s whole-dish sweep, its two nulls and its class rule, rerun on its own per-pixel ratio maps with only the region value changed; the same code with GENIE’s value reproduces GENIE’s record to 1.5 × 10⁻⁸. (c) Pearson r between a region and its ring against separation, the mean of the four dishes with a band of ± 1 SD, under GENIE’s value (purple, filled) and under the mean of the pixel ratios (orange, open), each with its within-field null (dotted). (d) The excess of r over the within-field null, mean ± 1 SD; the mean curve reaches the null at 32 µm under both values. (e) The first-ring test per dish: the observed r, the within-field null and the excess under both values (filled, GENIE’s; open, the mean of the pixel ratios), with the mean of the four dishes as a bar; the paired t test on the excess gives t = 5.04, p = 0.015 under GENIE’s value and t = 5.00, p = 0.015 under the mean of the pixel ratios. (f) The ring partners of HIGH and LOW regions (above or below the dish mean by 2 SD) minus the dish mean, in units of ΔΨm, against separation, under both values (solid, GENIE’s; dashed, the mean of the pixel ratios), the mean over the dishes that have the class.

## Notes

https://doi.org/10.5281/zenodo.23245979

